# Expression and secretion of circular RNAs in the parasitic nematode, *Ascaris suum*

**DOI:** 10.1101/2022.02.08.479594

**Authors:** Sarah J. Minkler, Hannah J. Loghry-Jansen, Noelle A. Sondjaja, Michael J. Kimber

## Abstract

Circular RNAs (circRNAs) are a recently identified RNA species with emerging functional roles as microRNA (miRNA) and protein sponges, regulators of gene transcription and translation, and as modulators of fundamental biological processes including immunoregulation. circRNAs have been found in a variety of species including plants, animals, and model genetic organisms such as the free-living nematode *Caenorhabditis elegans*. Relevant to this study, circRNAs have recently been described in the parasitic nematode, *Haemonchus contortus*, suggesting they may have functionally important roles in parasites. Given this involvement in regulating biological processes, a better understanding of their role in parasites could be leveraged for future control efforts. Here, we report the use of next-generation sequencing to identify 1,997 distinct circRNAs expressed in adult female stages of the gastrointestinal parasitic nematode, *Ascaris suum*. We describe spatial expression in the ovaries and body wall muscle, and also report circRNA presence in extracellular vesicles (EVs) secreted by the parasite into the external environment. Further, we used an *in-silico* approach to predict that a subset of *Ascaris* circRNAs bind both endogenous parasite miRNAs as well as human host miRNAs, suggesting they could be functional as both exogenous and endogenous miRNA sponges to alter gene expression. There was not a strong correlation between *Ascaris* circRNA length endogenous miRNA interactions, indicating *Ascaris* circRNAs are enriched for *Ascaris* miRNA binding sites, but that human miRNAs were predicted form a more thermodynamically stable bond with *Ascaris* circRNAs. These results suggest that secreted circRNAs could be interacting with host miRNAs at the host-parasite interface and influencing host gene transcription. Lastly, although we previously found that therapeutically relevant concentrations of the anthelmintic drug ivermectin inhibited EV release from parasitic nematodes, we did not observe a direct effect on *Ascaris* circRNAs expression or secretion.

## Introduction

Circular RNAs (circRNAs) are a species of long, noncoding RNA that do not contain an open 5’ or 3’ end but instead form a circular structure that is more stable than linear RNA species (Enuka et al., 2016). The majority of circRNAs are approximately 1500 nucleotides (nt) or less and have a median length of 550 nt (Zheng et al., 2016; Ding et al., 2018). circRNAs were first discovered through electron microscopy imaging of HeLa cells, CV-1 cells (monkey kidney cell line), and Chinese hamster ovary cells (Hsu and Coca-Prados, 1979) and initially thought to be the product of nontraditional splicing, forming “scrambled exons” with no real function or significance (Nigro et. al, 1991). With advancements in the sensitivity of high throughput sequencing and data analysis pipelines, the complexity of the circRNA complement has been recognized, validated, and shown to be functionally active in a variety of species including humans (Memczak et. al, 2013; Guo et. al 2014), mice (Memczak et. al, 2013), insects (Westholm et al., 2014), plants (Zhang et al., 2020), fungi (Shao et al. 2019), and germane to the current study, the model nematode *Caenorhabditis elegans* (Memczak et al., 2013; Ivanov et al., 2015; Cortés-López et al., 2018). The recognition that circRNAs are expressed in *C. elegans* has recently seeded their discovery in parasitic nematodes, specifically, the small ruminant gastrointestinal parasitic nematode, *Haemonchus contortus* (Zhou et al., 2021).

The biogenesis of circRNAs is summarized in Figure 1. Exonic (containing only exons) and intergenic (containing both exons and introns) circRNAs are formed when pre-mRNA transcripts undergo a back-splicing event where a downstream splice donor site attacks an upstream splice acceptor site (Memczak et al., 2013). These splice sites are brought together through intron looping that is facilitated by inverted repeat base pairing (Ivanov et al., 2015) or via pairing of RNA-binding proteins (RBPs) (Conn et al., 2015; Errichelli et al., 2017). Distinct from this process, intronic circRNAs (containing intronic RNA only) are formed from lariat precursors during linear splicing that evaded debranching and remains in a circular structure to avoid degradation (Kristensen et al., 2019). A fundamental function of circRNAs is the regulation of gene expression, which is accomplished through multiple pathways. The most recognized is that circRNAs act as miRNA sponges, binding multiple miRNAs and influencing gene expression by reducing miRNA bioavailability. This property of miRNA binding was first discovered in mice by Hansen et al. (2013) who found that CDR1as could bind murine miRNAs and modify miRNA biological functions as a result. Numerous miRNA binding sites have also been found in *Drosophila* circRNAs (Westholm et al., 2014) in support of this role. In addition, circRNAs can also promote gene transcription through interactions with RNA polymerase II and U1 snRNP in the promoter region of a gene (Li Z. et al., 2015), and in some instances, circRNAs can also be translated into proteins but the function of circRNA translated proteins remains largely unexplored (Pamudurti et al., 2017).

**Figure 1.**
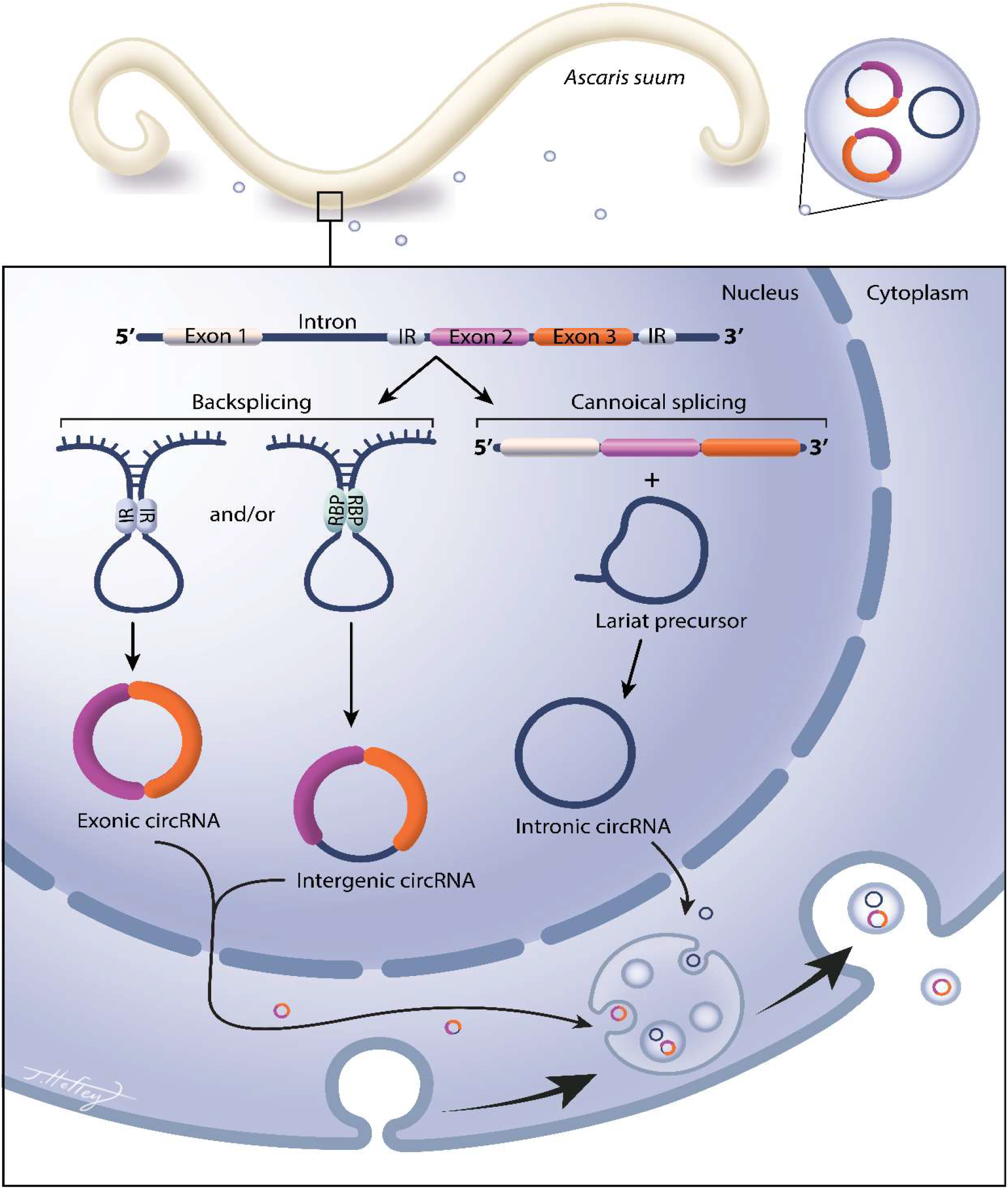
Circular RNAs are expressed in Ascaris suum and secreted into the host environment in extracellular vesicles. Circular RNA (circRNA) are covalently closed circular RNA rings, with no open 5’ or 3’ ends. They do not contain a polyA tail or a 5’ cap and are extremely stable and less prone to degradation as compared to linear RNA species. There are three distinctly recognized circRNA species; exonic, intergenic, and intronic circRNAs (Kristensen et. al., 2019). Exonic circRNAs contain only exons, while intergenic circRNAs contain both exons and introns. Intronic circRNAs contain only introns. Exonic and intergenic circRNAs are generated from a back splicing event where inverted repeats (IR) or RNA binding proteins (RBP) form a semi-closed covalent ring, allowing for downstream splice donor site to attack upstream slice acceptor site, forming the close circRNA structure. Intronic circRNAs are formed from lariat precursor molecules during linear splicing. circRNAs can leave the nucleus and become packaged into parasitic EV cargo for secretion, but this mechanism remains elusive.

Currently, there is a sparse amount of data on the expression of circRNAs in nematodes. As regulators of biological processes, a description of circRNA complement and function in parasites is warranted as they may have value from a control perspective. circRNAs have been identified in *C. elegans* (Memczak et al., 2013; Ivanov et al., 2015; Cortés-López et al., 2018) with the first descriptive study of a nematode circRNA complement based on *H. contortus* recently emerging (Zhou et al., 2021). These manuscripts focus on the presence of circRNAs in these two clade V nematode species but do not give much insight into the functional significance of circRNAs in worms. Here we describe the spatial expression of circRNAs in the clade III nematode *Ascaris suum. A. suum* is a large gastrointestinal parasite that primarily infects swine but has also been shown to infect humans. There are studies suggesting that *A. suum* and *Ascaris lumbricoides*, a human gastrointestinal nematode, are the same species due to cross infections between humans and pigs (Leles et al., 2012) and similarities in nucleic acid profiles (Shao et al., 2014; Nejsum et al., 2005). Infections with *A. suum* in pigs, leads to decreased farming productivity, and carries negative economic impacts including reduced animal growth, losses of meat product from contamination, treatment costs, and co-infections with other pathogens (Thamsborg et al. 2013). In humans, 807 million -1.2 billion people are infected with *Ascaris* worldwide (Centers for Disease Control, 2020). Infections with *Ascaris* can lead to gastrointestinal obstructions, anemia, diarrhea, hepatobiliary, and pancreatic syndromes. There is a disproportionate number of infections occurring in children, which can lead to malnutrition and cognitive impairment, leading to a loss of education (Bethony et al., 2006).

circRNAs are known to be secreted into the extracellular environment via extracellular vesicles (EVs) (Lasda et al., 2016), but have not been shown to be secreted in parasitic nematode EVs. Our laboratory and others, have previously shown that parasitic nematodes secrete EVs and that these EVs contain small RNA species (Zamanian et al., 2015; Buck et al., 2014; Hansen E. et al., 2019; Gu et al., 2017) but the presence of circRNAs has not been demonstrated in parasitic nematode EVs to date. We hypothesized that *A. suum* expresses endogenous circRNAs that may function as miRNA sponges. Further, that a cohort of these circRNAs are secreted and that these secreted circRNAs could interact with host miRNAs to have an impact at the host-parasite interface. To investigate these hypotheses, we collected EVs from *A. suum* adult female parasites and used next-generation sequencing to describe the endogenous circRNA complement. We then tested for the presence of secreted circRNAs within *A. suum* EVs. We found that circRNAs are expressed in body wall muscle and ovarian tissue and whilst some tissue-specific expression patterns were observed, our data indicate circRNAs have broadly distributed expression patterns in *Ascaris*. Select circRNAs were also found to be secreted in EVs and this is the first study to describe this mechanism in parasitic helminths. Predicted binding of both endogenous and exogenous circRNAs to host and *Ascaris* miRNAs led to the hypothesis that circRNAs are functioning as miRNAs sponges both endogenously and exogenously. These results suggest that circRNAs may function as miRNA sponges within *Ascaris* but, when secreted, may bind to host miRNAs and therefore influence host gene expression at the host-parasite interface.

## Materials and Methods

Healthy, adult female *Ascaris suum* were collected from an abattoir in Marshalltown, Iowa, USA. These parasites were thoroughly washed multiple times in *Ascaris* Ringer’s Solution (ARS) [(13.14 mM NaCl, 9.67 mM CaCl_2_, 7.83 mM MgCl_2_, 12.09 mM Tris, 99.96 mM sodium acetate, 19.64 mM KCl) with gentamycin (100 µg/ml), ciprofloxacin hydrochloride (20 µg/ml), penicillin (10,000 units/ml), streptomycin (10,000 µg/ml), and amphotericin B (25 µg/ml) at pH 7.87 (all Sigma Aldrich, St Louis, MO)] and then incubated at 35°C. The following day, parasites were checked visually for signs of bacterial or fungal contamination, and worms were discarded if present. Worms were cut along the ventral midline and the ovaries were gently removed for excision. Tissue was collected proximal to the bifurcation of the ovaries and rinsed with fresh ARS. Given the location of ovarian tissue collection, we acknowledge residual embryonated egg material may be present. Body wall tissue was collected directly anterior to the genital aperture and musculature was scraped from the underlying cuticle with a single-edge razor blade. Approximately 200 mg of body wall muscle and ovary tissue samples were obtained in each sample isolation. Tissues were either used for immediate RNA extraction or stored at -80°C until use.

### 2.2 Drug Treatment

Individual worms were treated with 0.1 µM or 1 µM (final concentration) of ivermectin, diethylcarbamazine, or levamisole (all Sigma-Aldrich) for 24 hours in 100 ml culture media in sterile 250 ml Erlenmeyer flasks. Drug concentrations were prepared from stock solutions dissolved in dimethyl sulfoxide (DMSO, Sigma-Aldrich). Conditioned media from drug treated and DMSO vehicle control worms was collected after the 24-hour time period and retained for downstream analysis. Body wall muscle and ovarian tissue samples were collected from these parasites as described for immediate RNA extraction or storage at -80°C until use.

### 2.3 EV Isolation and Quantification

EVs were collected as previously described using differential ultracentrifugation (Loghry et al, 2020; Harischandra et al., 2018; Zamanian et al., 2015). Media was filtered through 0.2 µm PVDF vacuum filters (Sigma-Aldrich) and centrifuged at 120,000 x g for 90 minutes at 4°C. The supernatant was decanted, and pellets were filtered through a PVDF 0.2 µm syringe filter (GE Healthcare, Chicago, IL) and centrifuged further at 186,000 x g for two hours at 4°C. EV samples were then resuspended to 500 µl in dPBS (Thermo Fisher Scientific, Waltman MA) and stored at -80°C until use.

EV quantification and size determination were performed using nanoparticle tracking analysis (NTA; Nano-Sight LM10, Malvern Instruments, Malvern, UK). EV imaging was performed using transmission electron microscopy. A 2 µL aliquot of isolated EV preparation was placed onto a carbon film grid for 1 minute. The drop was wicked to a thin film and 2 µL of uranyl acetate (2% w/v final concentration) was immediately applied for 30 seconds, wicked, and allowed to dry. Images were taken using a 200kV JEOL 2100 scanning and transmission electron microscope (Japan Electron Optics Laboratories, LLC, Peabody, MA) with a Gatan OneView camera (Gatan, Inc. Pleasanton, CA).

### 2.4 Circular RNA Isolation

Total RNA was extracted from adult female *A. suum* body wall and ovary tissues using a two-step process. First, worm tissues were homogenized in TRIzol reagent (Life Technologies, Carlsbad, CA) according to manufacturer recommendations. Following RNA isolation in TRIzol, total RNA was further purified using the miRNeasy Mini kit (QIAGEN, Hilden, Germany), following the manufacturer’s protocol. The total RNA generated was assessed for purity and quantified using a NanoVue spectrophotometer (General Electric, Boston, MA). Similarly, total RNA was extracted from EV enriched samples isolated from conditioned media using the miRNeasy Micro kit (QIAGEN), again following the manufacturer’s protocol. Total RNA from EV isolation supernatants (i.e. EV-depleted media) was extracted using Zymo ZR urine RNA isolation kit (Zymo Research, Irvine, CA), following manufacturer’s instructions. All linear RNA was subsequently prepared from each of these total RNA preparations by RNase R digestion (RNase R was provided by the Singh Laboratory, Iowa State University). 10 Units of RNase R used for each reaction along with 2µg total RNA. Reactions were incubated at 37°C for 45 minutes followed by heat inactivation for 65°C for 20 minutes. circRNA was then stored at -80°C until use. circRNA samples were sent to LC sciences for circRNA sequencing or transcribed into cDNA for qPCR validation.

### 2.5 CircRNA-seq Library Preparation

CircRNA quality was assessed with a Bioanalyzer 2100 and RNA 6000 Nano LabChip Kit (Agilent, CA, USA), allowing a minimum RNA integrity number (RIN) of 7 (Schroeder et al., 2006) before fragmentation using NEBNext® Magnesium RNA Fragmentation Module (NEB, Ipswich, MA) into short fragments using divalent cations under high temperature. The cleaved RNA fragments were then reverse-transcribed to create the cDNAs using SuperScript™ II Reverse Transcriptase (Thermo Fisher Scientific), which were next used to synthesize U-labeled second-stranded DNAs with *E. coli* DNA polymerase I (NEB), RNase H (NEB) and dUTP Solution (Thermo Fisher). An A-base was added to the blunt ends of each strand, preparing them for ligation to the indexed adapters. Each adapter contains a T-base overhang for ligating the adapter to the A-tailed fragmented DNA. Single- or dual-index adapters were ligated to the fragments, and size selection (300-600bp) performed with AMPureXP beads (Beckman Coulter, Brea, CA). After the heat-labile UDG enzyme (NEB) treatment of the U-labeled second-stranded DNAs, the ligated products were amplified with PCR by the following conditions: initial denaturation at 95°C for 3 min; 8 cycles of denaturation at 98°C for 15 sec, annealing at 60°C for 15 sec, and extension at 72°C for 30 sec; and then final extension at 72°C for 5 min. The average insert size for the final cDNA library was 300 ±50 bp. Finally, 2×150bp paired-end sequencing (PE150) was performed on an Illumina Novaseq™ 6000 (Illumina) following the vendor’s recommended protocol.

### 2.6 CircRNA assembly

Cutadapt (Martin, 2011) and custom perl scripts were used to remove adaptors, low quality bases and undetermined bases, followed by quality assessments with FastQC (Andrews, 2010). Bowtie2 (Langmead & Salzberg, 2012) and Tophat2 (Kim et al., 2013) were used to map reads to the genome of *A. suum* (Wang et al., 2017) (Accession number: PRJNA62057; AGO1), with remaining unmapped reads remapped to the genome using Tophat-fusion (Kim & Salzberg, 2011). Dual de-novo assemblies of circular RNAs were performed with CIRCExplorer (Zhang et al., 2016; Zhang et al., 2014), one with Bowtie2 and Tophat2 mapped reads and another with Tophat-fusion back-spliced reads. Since samples were not prepared simultaneously, the distinct circRNA assemblies were concatenated and filtered for uniqueness with duplicated sequences being removed.

### 2.7 Analysis of circRNA-seq expression data

Sequenced RNA reads (SRR) (SRR15295818-SRR15295823) were aligned to circRNAs from both body wall and ovary, and to the *A. suum* genome (PRJNA62057), to reduce bias by avoiding creating mapping reads that are artifacts from flawed methodology (Wang et al., 2017) using Hisat 2.2.0 (Kim et al., 2016). Samtools 1.10 (Li et al., 2009) was used to convert sam alignments to bam alignment files to map RNA sequencing reads to the *A. suum* genome and the three circRNA samples. Mapping statistics were assessed using Picard 2.17.0 (Intitute, 2019). Read counts were taken using featureCounts from the Subread package 1.6.0 (Liao et al., 2013). Differential expression was assessed using DESEQ2 1.20.0 (Love et al., 2014), with both unique and multiple mapping reads considered in separate comparisons. Differentially expressed circRNAs were subjected to GO and KEGG enrichment analyses using clusterProfiler (Yu et al., 2012) and Ontologizer (Bauer et al., 2008). The number of exonic, intergenic, and intronic circRNAs were counted from sequencing data and visualized using GraphPad Prism version 9.2.0 (GraphPad Software Inc., San Diego, CA). Notes and scripts used to produce expression analysis are available in Supplementary Figure 6. Raw data and circRNA sequences can be viewed using bio-project number PRJNA750737 with SRA numbers, SRR15295818 - SRR15295823 (Supplemental Figure 7).

### 2.8 qRT-PCR CircRNA Validation

Validation that circRNAs identified using circRNA-seq are expressed in *Ascaris* tissue or EV enriched samples was performed using quantitative real-time PCR (qRT-PCR). RNase R treated RNA (generated as described above) was reversed-transcribed to cDNA with random hexamers using Invitrogen SuperScript III First Strand Synthesis System (Thermo Fisher Scientific) according to the manufacturer’s instructions. A total of 10 sets of divergent primers were designed to specific circRNA back-splice junction and qRT-PCR was performed using Power Up Sybr Green Master Mix according to the manufacturer’s protocol (Thermo Fisher Scientific). Conditions for qRT-PCR were as follows, 2 minutes at 50°C, 2 minutes at 95°C, then 40 cycles of 15 seconds at 95°C, 15 seconds at 55-60°C, and 1 minute at 72°C. CircRNA abundance was quantified by extrapolating qRTPCR C_T_ values from exogenous (spiked in) *Homo sapiens* actin alpha cardiac muscle 1 RNA (GenBank accession number: NM_005159). This spike in RNA was generated using MEGAScript T7 transcription kit (Thermo Fisher Scientific) following manufacturer’s protocols corresponding to 517 bp – 1,165 bp of the published transcript (primer sequences are detailed in Supplemental Figure 2). The spike in RNA standard curve analysis was created using Graph Pad Prism version 9.2.0 (GraphPad Software Inc.) using a nonlinear second-degree polynomial, least squares fit. X values were interpolated to quantify the concentration of circRNA in each tissue type and EVs.

**Figure 2.**
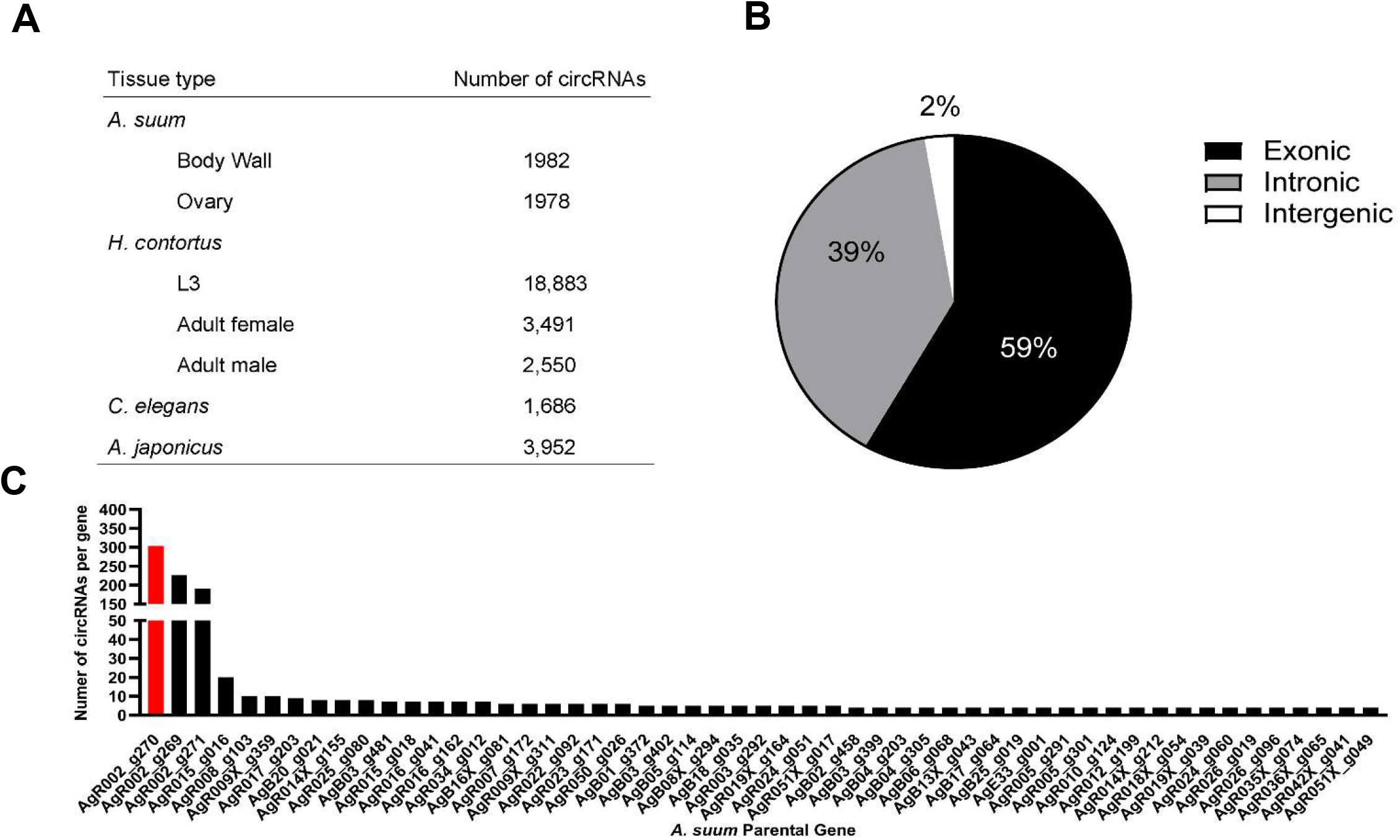
Characteristics of circRNAs identified in *A. suum* through next-generation sequencing. **(A)** 1,982 *A. suum* circRNAs were identified in body wall tissue and 1,978 circRNAs were identified in ovary tissue. The total amount of *A. suum* circRNAs identified in both tissues was 1,997. In H. contortus there was a total of 20,073 circRNA were discovered with the highest amount being found in L3 (18,883). In C. elegans, there was a total of 1686 exonic circRNAs discovered over a variety of life-stages, but there are no reports on the amount of intronic circRNAs found in this nematode. 3,952 circRNAs were identified in *A. japonicus* adults. **(B)** There are three recognized types of circRNAs that can be generated. Exonic circRNAs are derived from only exonic regions of mRNA, intergenic circRNAs contain both exons and introns, and intronic circRNAs are formed from lariat precursors and only contain intronic regions. **(C)** There were a number of *A. suum* parental genes that had more than one circRNA derived from it. AgR002_g270 had the highest amount of circRNAs (303) and is colored in red. Only genes with over 4 circRNAs were included in this graph.

### 2.9 CircRNA-miRNA interactions

Two complementary programs were used to predict miRNAs and circRNA interactions: miRanda (https://github.com/hobywan/miranda) and Targetscan (Agarwal et al., 2018) using the identified *Ascaris* circRNA sequences and miRNA datasets for human and *A. suum* downloaded from miRbase.org (Release 22.1)(Kozomara et al., 2010). Within these programs, a higher miRanda free energy (200-140) and lower Targetscan score (−0.13-0) were used to assign interaction confidence. The number of *Ascaris* and human miRNA interactions for each *Ascaris* circRNA was totaled and visualized using GraphPad Prism (GraphPad Software Inc.). Similarly, *Ascaris* and human miRNAs with high numbers of predicted *Ascaris* circRNA binding partners were also collated and visualized using GraphPad Prism (GraphPad Software Inc.).

### 2.10 Statistical Analysis

IVM treated tissues and EV circRNA expression levels were calculated from qRT-PCR CT values using 2^−ΔΔCq^ (Livak & Schmittgen, 2001). Following fold change analysis, the data was log(2) transformed using GraphPad Prism (GraphPad Software Inc.). To compare circRNA concentrations between treatments and samples, a two-way ANOVA with multiple comparisons was used with a p-value less than 0.05 being significant in GraphPad Prism (GraphPad Software Inc.). Each N number represented a new batch of worms, with an individual no treatment control for each batch.

## Results

### 3.1 A. suum circRNA Complement

To examine the presence of circRNA in *A. suum* tissues, a total of six independently prepared samples (three ovary, three body wall) were used for circRNA sequencing. After the removal of redundant or duplicated circRNAs, we identified a total of 1,982 circRNAs in body wall tissue and 1,978 circRNAs in ovary tissue, for a total of 1,997 unique and distinct circRNAs (Figure 2A). There was a significant number of circRNAs shared between the two tissue types (1,963) and only 34 circRNAs were identified through circRNA-seq analysis as having tissue-specific expression. 15 circRNAs were identified only in the ovary-enriched samples and 19 were body wall specific (Figure 4C). Tissue specific circRNAs are listed in Supplementary Figure 1. If these circRNAs are functional, such a general spatial distribution pattern suggests either that the majority of *A. suum* circRNAs are linked to the regulation of broadly conserved transcriptional processes, or that any functional specificity is driven by a more restricted temporal or spatial expression of interacting partners rather than the circRNA itself. Raw data and circRNA sequences can be viewed using bio-project number PRJNA750737 with SRA numbers, SRR15295818 - SRR15295823 (Supplemental Figure 7).

**Figure 3.**
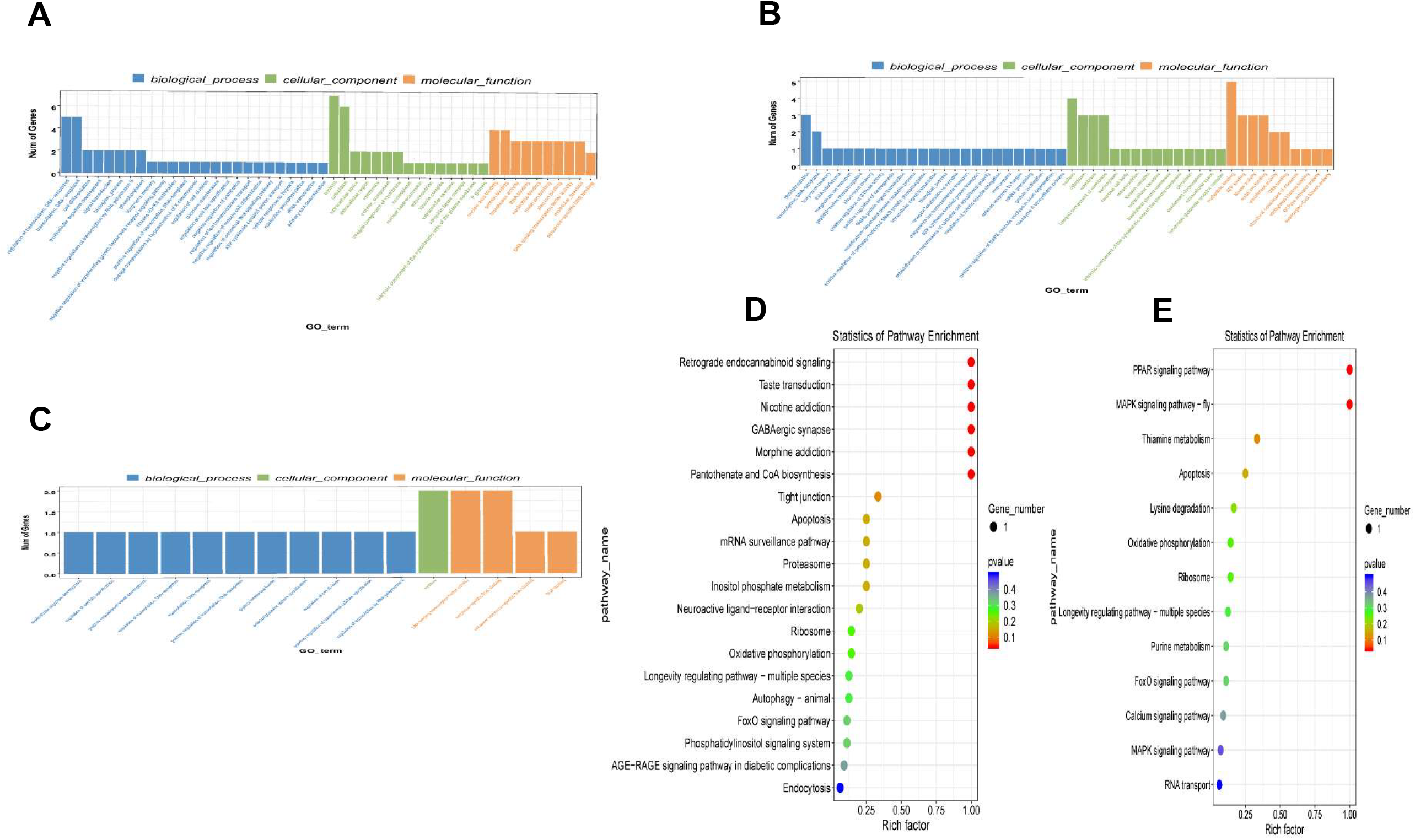
GO and KEGG term analysis for differentially expressed circRNAs in *A. suum* ovary and body wall tissues. A total of 6 samples were sent for sequencing (3 body wall and 3 ovary tissue samples). Each sample was compared against the other two samples of the different tissue. Significant GO and KEGG terms were calculated using the hypergeometric equation. **(A-C)** Individual GO term graphs of ovary vs. body wall tissue from each individual sample that was submitted to sequencing. Different color bars are representing processes (biological process, cellular function, molecular function). Number of differentially expressed genes on y-axis. **(D-E)** Enriched KEGG terms for each sample submitted to sequencing. One sample did not have any significant differentially expressed KEGG terms. X-axis represents rich factor. Rich factor is the ratio of the number of differentially expressed genes annotated in a pathway. The color and size of each bubble represent p-value and the number of genes enriched in a pathway.

**Figure 4.**
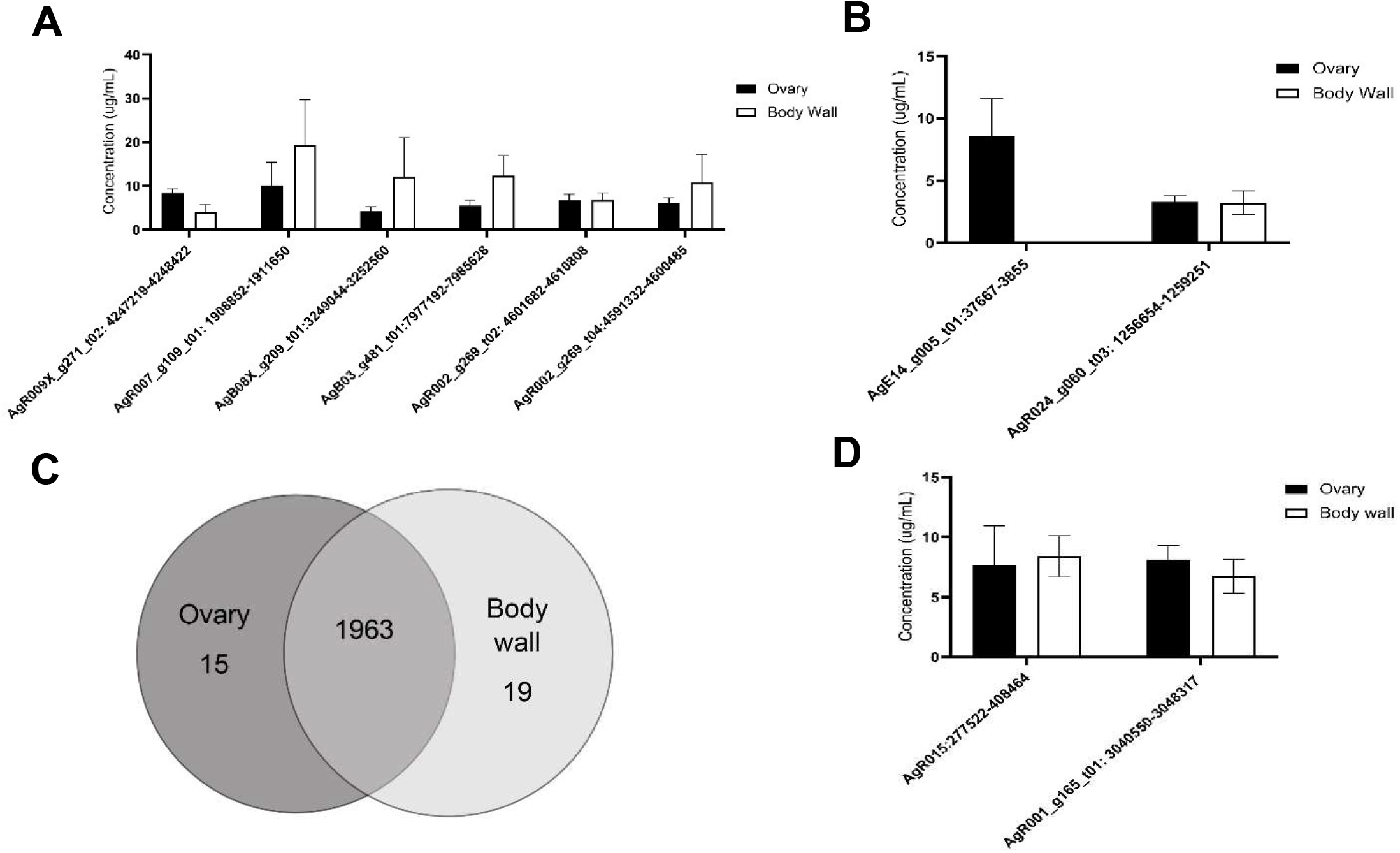
RT-qPCR validation of ten individual circRNAs in *A. suum* ovary and body wall tissues. circRNAs identified through sequencing were validated for tissue expression using RT-qPCR analysis. CT values were normalized with standard curve analysis using spike in RNA to calculate concentration (µg/mL) for each circRNA. N=4 (minimum), mean ± SEM, p-value less than 0.05 being significant. **(A)** circRNA concentrations of six select circRNAs in ovary and body wall tissues. These circRNAs were prioritized because of their robust expression is our circRNA sequencing data across both body wall and ovary tissue samples. There was no statistical significance between ovary and body wall tissues for any of the six circRNAs examined. **(B)** The spatial expression of two circRNAs with tissue specific distribution in our circRNA sequencing datasets were analyzed using RT-qPCR. AgE14_g005_t01:37667-3855 was expected to be expressed in ovary tissue only and this was validated. Although AgR024_g060_t03:1256654-1259251 was expected to be expressed in body wall tissue only based on sequencing data, PCR results suggest a broader spatial distribution pattern (N=3 minimum). **(C)** A total of 1963 circRNAs were common to both tissue types, 15 circRNAs were found only in the ovary samples while 19 were specific to the body wall preparations. **(D)** A number of putative circRNAs of atypically large size (over 5 kb) were identified in our sequencing data. Expression of two such circRNAs was validated by qPCR. Amplification using primers spanning back-splice junctions indicates these large RNA molecules are circRNAs.

Of the total 1,997 circRNAs identified, 1,178 (59%) were exonic, 779 (39%) were intronic and 40 (2%) were intergenic (Figure 2B). Whilst there is a lack of data distinguishing the functional importance of exonic from intronic circRNAs, there is the potential for exonic circRNAs to be translated into proteins (Legnini et al., 2017), and those translated proteins could have important biological roles.

The number of exons per circRNAs was calculated and on average, circRNAs contained approximately three exons and 83% of circRNAs were composed of multiple exons (two or more). Interestingly, 37% (752) of circRNAs seem to be derived from chromosome AgR001, while gene AgR002_g270 had the most derived circRNAs, a remarkable 15% of the total (303) (Figure 2C). AgR002_g270 does not have a known function or any identified ortholog or paralogs associated with it, but blastx analysis (https://blast.ncbi.nlm.nih.gov) of the AgR002_g270 coding sequencing returns ribosomal proteins from various nematode species, including *A. lumbricoides, Ascaridia galli, Toxocara cati*, and *Baylisascaris procyonics*.

### 3.2 GO and KEGG analysis of circRNA parental genes

CircRNA have the potential to encode proteins (Legnini et al., 2017) so understanding the function of parental genes from which *Ascaris* circRNA derive could provide information about circRNA function. GO and KEGG annotation analyses were conducted to predict possible determine functions of parental genes (Figure 3). Significant GO and KEGG terms were calculated by hypergeometric equation (Erdélyi et al., 1953) and terms with p-values less than 0.05 were defined as significant. Significant GO terms were divided into three groups, biological process, cellular component, and molecular function. 49 GO terms were involved in the biological process category for all three replicates (duplicates were excluded). The most enriched GO terms in the biological process category included “regulation of transcription; DNA templated” (GO:0006355), “transcription, DNA templated” (GO:0006351); and “phosphorylation” (GO:0016310) (Figure 3A-C). For cellular component, 41 individual GO terms were significantly enriched. They mainly consisted of “nucleus” (GO:0005634) and “cytoplasm” (GO:0005737). Molecular function had one enriched GO term, “nucleotide binding” (GO:0000166) (Figures 3A-C).

KEGG pathway analysis was also carried out to determine further significant pathways of circRNA parental genes and identify enriched pathways. There was a total of three different KEGG comparison groups, one for each of the ovary and body wall samples that were submitted from the same adult female worm. Of the three different comparisons that were analyzed for differential expression, one sample comparison did not have any significant differentially expressed KEGG terms and is not included in this analysis. The most enriched KEGG pathways for differential expression between ovary-enriched and body wall tissues include PPAR signaling pathway, MAPK signaling pathway, and retrograde endocannabinoid signaling (Figure 3D-E). Ten out of twenty-eight of differentially expressed KEGG pathways were involved in “signaling”, suggesting that circRNA parental genes are involved with signaling and signal transduction pathways and other important cellular processes to support worm health. If *A. suum* circRNAs are translated and yield functional proteins, this GO and KEGG term analysis points to possible functions that circRNAs could be performing in the worm, based on parental gene functions.

### 3.3 qRT-PCR tissue validation of A. suum circRNA expression in ovary and body wall tissue

We used qRT-PCR to confirm and validate the expression of six individual circRNAs identified in *A. suum* samples using circRNA-seq. Six initial circRNAs were selected due their high-count numbers from sequencing data in both ovary and body wall tissues. Divergent primers spanning back-splice junction sites were designed for each circRNA and can be viewed in Supplemental Figure 2. Using an RNA spike in approach allowed us to calculate circRNA concentration levels using a standard curve, and as expected we did not observe any statistical significance in the abundance of individual circRNAs between the two tissue types (N=5) (Figure 4A). This data validated the circRNA-seq approach as a means to broadly describe circRNA expression but further qRT-PCR analyses of differentially expressed circRNAs underscored the importance of verifying the circRNA-seq data with secondary methods (Figure 4B-C). The circRNA-seq datasets identified AgE14_g005_t01:37667-3855 as specifically expressed in the ovary-enriched samples and AgR024_g060_t03: 1256654-1259251 in the body wall samples. A full list of tissue specific circRNA expression for both ovary and body wall can be viewed in Supplementary Figure 1. Although our qRT-PCR data confirmed that AgE14_g005_t01:37667-3855 was indeed localized to ovary tissue, AgR024_g060_t03: 1256654-1259251 was found to be expressed in both tissue types (Figure 4B). This result could be due to the ability of qRT-PCR to amplify partially degraded transcripts and has been observed in other species (Westholm et al, 2014; Zhao et al., 2019) or could be accounted for by contamination of the ovary-enriched sample with body wall circRNA during dissection or from cross-contamination of RNA samples. It underscores the need for increased depth of circRNA sequencing married with additional validation measures to confirm spatial localization of circRNAs.

Further validation of select circRNA expression was performed, specifically, of AgR015: 277523-408464 and AgR001_g15_t01: 3040550-3048317. These circRNAs were prioritized due to their large size: AgR015: 277523-408464 was 130,941 nt long and AgR001_g15_t01: 3040550-3048317 was 7,767 nt long. We were able to confirm that these two RNA molecules are circRNAs through qPCR validation (Figure 4D) by creating primers specific to their back-splice junction. This approach demonstrates the RNAs form a circular structure and are not spurious background artifacts or RNA molecules residual from RNase R digestion even though they are larger in size than typical circRNAs (approximately 500-600 nt)(Ding et al., 2018).

### 3.4 circRNAs are secreted in extracellular vesicles, but circRNA expression is not grossly affected by ivermectin treatment

circRNAs have been found to be secreted from mammalian parental cells in extracellular vesicles (EVs) (Lasda et al., 2016). Many species of nematodes are known to secrete EVs (Buck et al., 2014; Zamanian et al., 2015; Harischandra et al., 2018; Hansen E. et al., 2019; Eichenberger, Talukder et al., 2018; Eichenberger, Ryan et al., 2018; Tzelos et al., 2016; Tritten et al., 2017; Shears et al., 2018) but the presence of circRNAs in those vesicles has not been explored. We hypothesized that *A. suum* EVs would contain circRNAs. To test this hypothesis, we used qRT-PCR to quantify the abundance of select circRNA transcripts in *A. suum* EVs using the spike-in approach as previously described. We first isolated, TEM imaged and performed nanoparticle tracking analysis (NTA) on *A. suum* EVs to confirm EV presence and structure (Figure 5A-B). Given previous data published by our laboratory on the inhibitory effect of ivermectin (IVM) on parasitic nematode EV secretion, we also examined whether IVM would inhibit circRNA secretion if present in the EVs. Parasites were cultured in the presence or absence of 0.1 µM (Figure 5D) and 1 µM (Supplemental Figure 3) IVM to model a therapeutically relevant dose. After 24 hours, parasite media was collected, and total EV RNA was extracted and used in RT-qPCR. We tested the presence of the same six circRNAs that were also tested for in tissue qRT-PCR validation (Figure 4A). Of these, two circRNAs (AgR007_g109_t01: 1908852-1911650 and AgB08X_g209_t01:3249044-3252560) could be consistently amplified from EV RNA samples (N=3) (Figure 5C). These data indicate that parasitic nematode circRNAs are secreted into the host milieu in EVs. We also looked for the presence of circRNAs in *A. suum* supernatants generated through EV isolation, representing non-EV mediated mechanisms of circRNA secretion. This approach yielded insufficient amounts of total RNA to conduct RT-qPCR analysis suggesting EVs may represent the primary mechanism of circRNA secretion from this worm.

**Figure 5.**
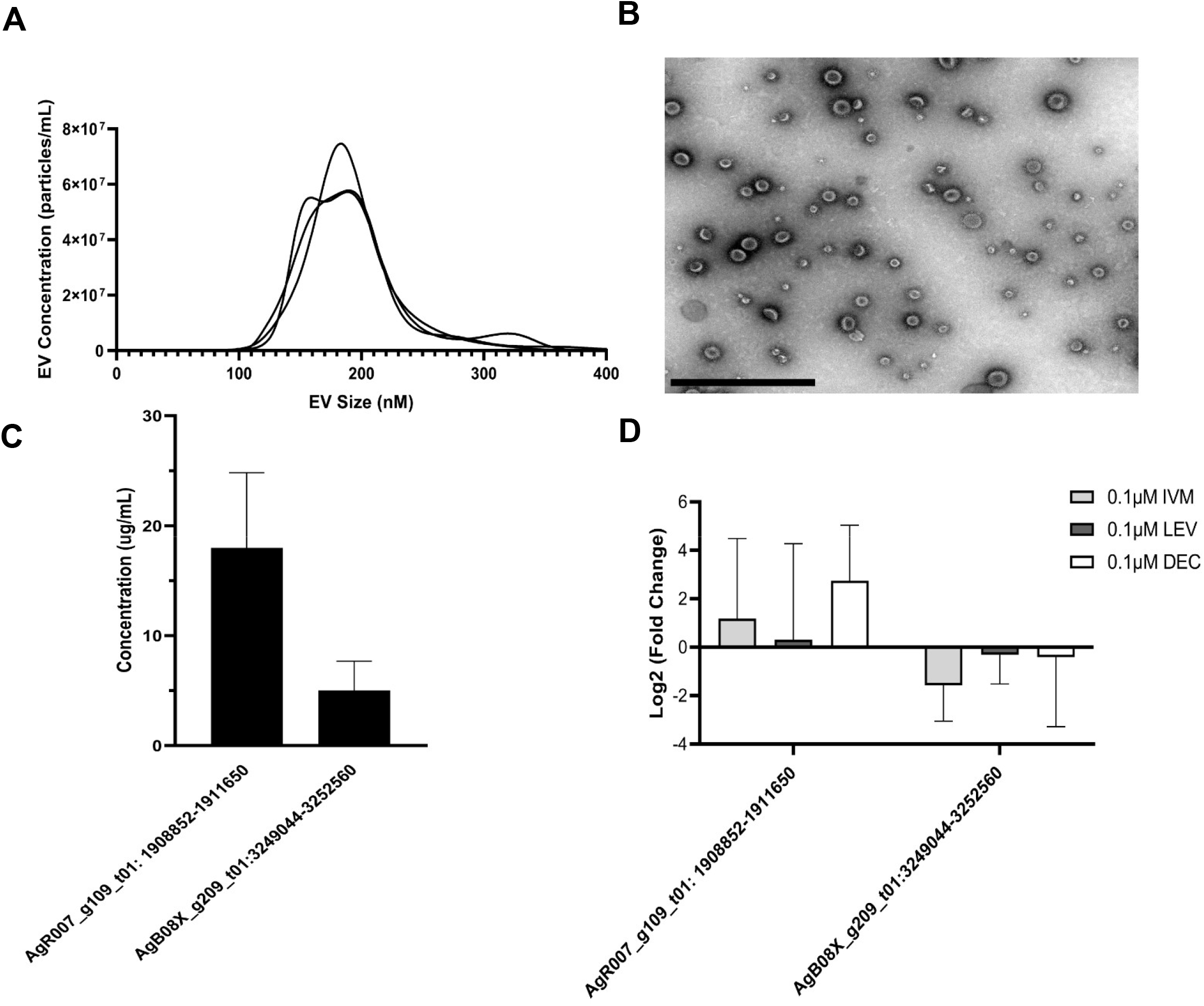
circRNA expression in parasitic extracellular vesicles after DMSO, IVM, DEC, and LEV treatment. **(A)** EVs were isolated using differential ultracentrifugation and NTA was used to determine EV size and concentration. EVs average around 200nM in size, which is similar to other reports on *A. suum* EVs (Loghry et al., 2020) **(B)** Extracellular vesicles (EVs) from A. suum adult female worms were collected through differential ultracentrifugation and imaged under TEM to validate presence and structure before determining circRNA expression in EVs. Scale bar represents 1000nm. qRT-PCR CT values were normalized to 40ng spike in RNA using 2^−ΔΔCq^ and log(2) transformation was also performed. N=3 (minimum), Mean ± SEM, p-value less than 0.05 being significant. **(C)** circRNA expression levels in untreated A. suum EVs. Concentration was calculated using a standard curve normalized to spike in RNA. Only two out of six circRNAs were detected in *A. suum* EVs with accurate reproducibility. **(D)** circRNA expression in EVs was unaffected by 24 h treatment of parental parasites with therapeutically relevant doses of the anthelmintic drugs ivermectin (IVM), diethylcarbamazine (DEC), or levamisole (LEV). circRNA expression in EVs was normalized to EVs secreted by untreated control worms normalization to control EVs.

When worms were treated with 0.1 µM (Figure 5B) or 1 µM (Supplemental Figure 4) IVM we did not observe any decrease in AgR007_g109_t01: 1908852-1911650 or AgB08X_g209_t01:3249044-3252560 abundance in isolated EVs (Figure 5B, N=3). This observation was perhaps surprising, given the strong and consistent evidence for an inhibitory effect of IVM on EV secretion in parasitic nematodes, including *Ascaris* (Loghry et al., 2020; Harischandra et al., 2018). This may point to other non-EV mediated routes of circRNA release from these worms. Other anthelmintic drugs are reported to have sporadic inhibitory effects on EV secretion by some life stages of filarial parasitic nematodes (Loghry et al., 2020). Therefore, we also looked at the effect of diethylcarbamazine (DEC) and levamisole (LEV) treatment on circRNA expression in *A. suum* EVs (Figure 5D, Supplemental Figure 3). Consistent with the IVM data, we did not see any inhibition in EV circRNA abundance, collectively indicating inhibition of circRNA secretion via EVs is not clearly associated with the mode of action of anthelmintic drugs.

To fully evaluate the effect of anthelmintic drug treatment on circRNA expression, we lastly examined whether IVM, DEC, or LEV had any effect on circRNA expression in *Ascaris* tissues using the same cohort of six prioritized circRNAs. Treatment of worms with 0.1 µM IVM (Figure 6) or 1µM (Supplementary Figure 4) did not alter expression of any tested circRNA in ovary tissues (N=5). Similarly, five of the six circRNAs in body wall tissue were unaffected by IVM treatment although AgR002_g269_t04:4591332-4600485 was significantly downregulated by 95% compared to control (p = 0.0071, N=4) in body wall tissue at 0.1µM (Figure 6), but not at 1µM (Supplementary Figure 4). Consistent with the results from IVM treated tissues, we did not observe any effect of DEC or LEV (Supplemental Figure 5) on circRNA expression in *A. suum* ovary or body tissues. Collectively our data do not support the hypothesis that anthelmintic drug mechanism of action involves a direct impingement of normal circRNA expression or secretion.

**Figure 6.**
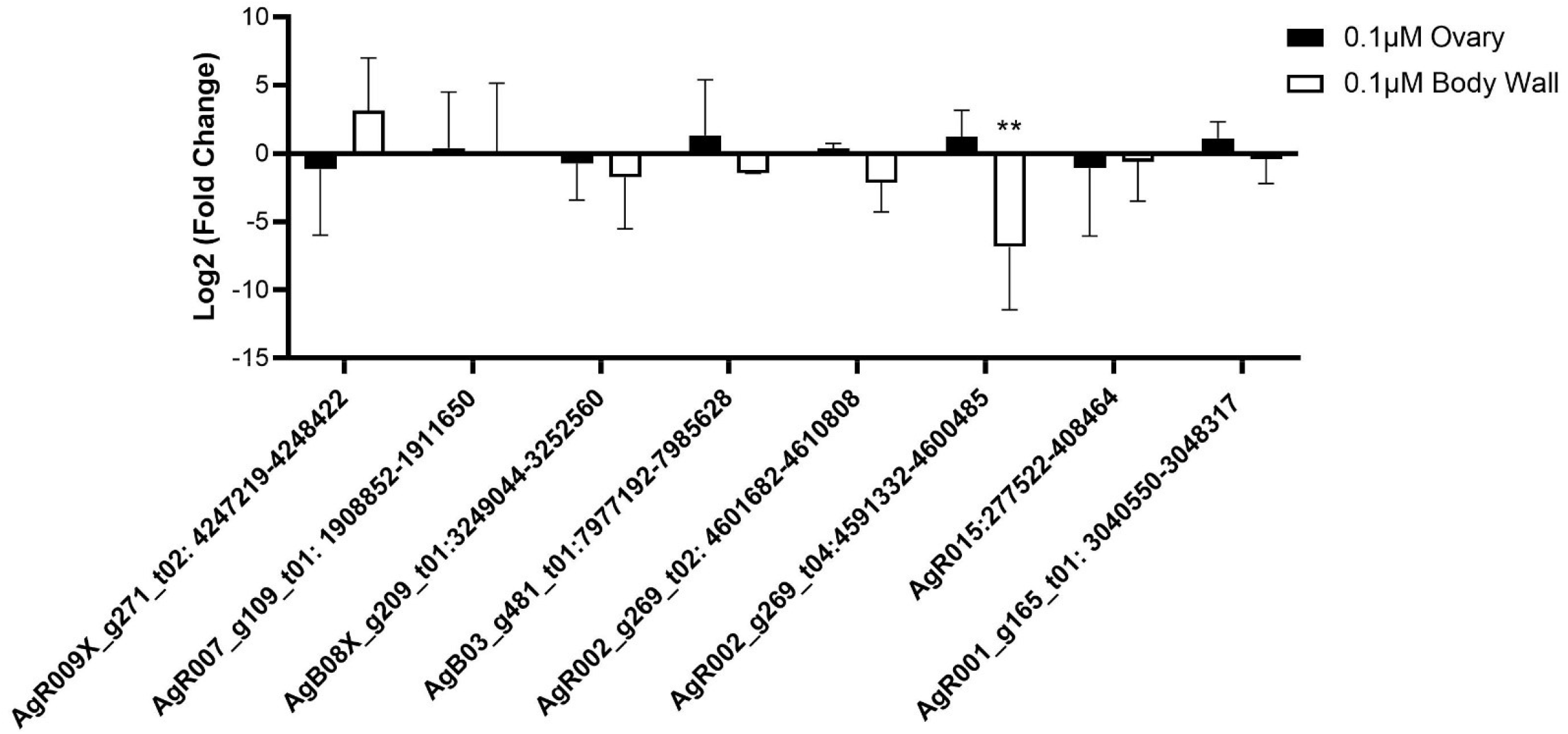
IVM has minimal effects on native circRNA expression in *A. suum* ovary and body wall tissues. qRT-PCR CT values were normalized to 40ng spike in RNA using 2^−ΔΔCq^ and log(2) transformation was also performed. N=4 (minimum), Mean ± SEM, p-value less than 0.05 being significant, *P<0.05, **P<0.01. circRNA expression levels in IVM treated ovary tissue after 24 hours incubation. No significant effect of IVM was observed in ovary tissue 0.1µM. IVM treated body wall was overall, not strongly affected by IVM treatment except for AgR002_g269_t04:4591332-4600485 at 0.1µM IVM. This was the only circRNA that was significantly different from control for both tissues and concentrations after IVM treatment.

### 3.6 Ascaris circRNAs are predicted to act as miRNA sponges

A well-established functional role for circRNAs is to impact gene expression by binging miRNAs, effectively acting as miRNA sponges. circRNAs can contain numerous binding sites for either individual or multiple miRNAs (Li F. et al., 2015; Capel et al., 1993; Zheng et al., 2016) For instance, murine CDR1as (ciRs-7) has 63 conserved binding sites for the miRNA mir-7 (Hansen T.B. et al., 2013), while circHIPK3 can sponge to nine different human miRNAs (Zheng et al., 2016). Here, we wanted to probe potential miRNA interactions with the *A. suum* circRNA dataset to support the hypothesis that *Ascaris* circRNAs can act as a miRNA sponges. Given the evidence that *Ascaris* circRNAs are secreted into the host via EVs, we used the miRanda algorithm (Enright et al., 2003) to predict interactions between *Ascaris* circRNAs and both endogenous *A. suum* miRNAs and host (human) miRNAs.

202 out of 1,997 *A. suum* circRNAs (∼10%) were predicted to interact with *A. suum* miRNAs (Figure 7A). The number of miRNA interactions per circRNA varied, but AgR015:277522-408464 was found to have 174 distinct miRNA interactions, being the highest number of miRNA interaction number. While AgR015:277522-408464, was predicted to bind 174 *A. suum* miRNAs, there were only 19 identified binding sites for this circRNA suggesting that different miRNAs can bind to the same sites on circRNAs. There were more predicted interactions between *Ascaris* circRNAs and human miRNAs at 398 (Figure 7B), with AgR015:277522-408464 having the highest number of predicted miRNA interactions (2308).

**Figure 7.**
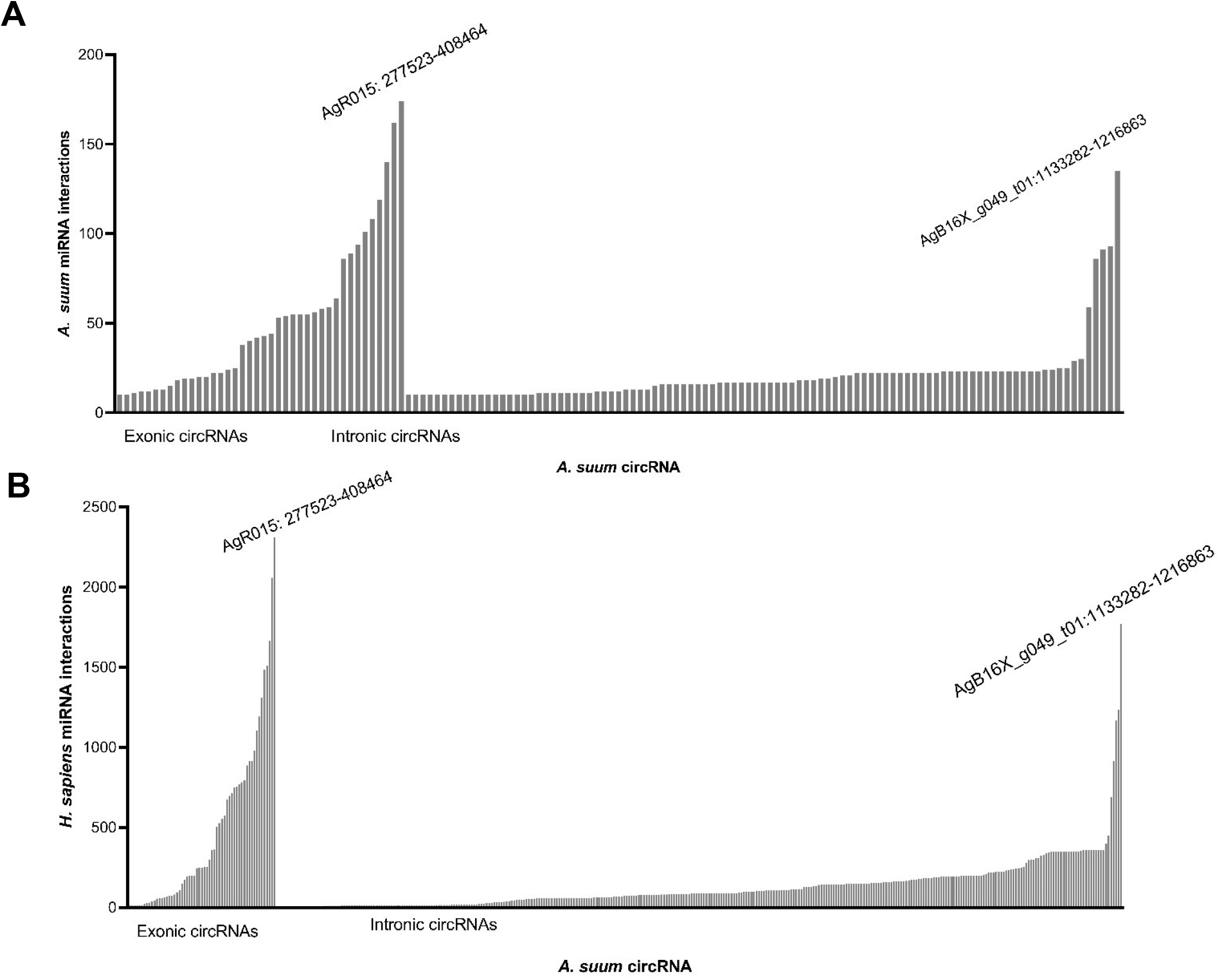
*A. suum* circRNAs interact with both endogenous and host miRNAs. Graph Pad Prism was used to graphical represent circRNA predicted interactions with worm and host miRNAs. circRNAs with under 10 miRNA interactions were excluded from these graphs. **(A)** circRNAs with predicted *A. suum* miRNA interactions. Exonic circRNAs had a larger number of A. suum miRNA interactions as compared to intronic circRNAs. Exonic circRNA, AgR015: 277523-408464 had the highest amount of predicted *A. suum* miRNA interactions at 174. Intronic circRNA AgB16X_g049_t01:1133282-1216863 had the most amount of predicted worm miRNA interactions (135). **B)** circRNAs with over 10 human miRNA interactions. circRNAs had more predicted human miRNA interactions as compared to worm miRNAs. *A. suum* exonic circRNAs again, had more interactions with human miRNAs as compared to intronic circRNAs, with AgR015: 277523-408464 having the most exonic circRNA interactions at 2,308. Intronic circRNA AgB16X_g049_t01:1133282-1216863 had the highest number of predicted human miRNA interactions at 1,769.

This disparity may reflect the greater number of annotated miRNAs in the human genome compared to that of *Ascaris*. Interestingly, for both human and *Ascaris* miRNAs, exonic circRNAs exhibited a higher number of predicted miRNA interactions than intronic circRNAs (Figure 7). Interactions between exonic and intronic circRNAs with Ascaris and host miRNAs is summarized in Table 1. The highest number of interactions between exonic circRNAs and *Ascaris* miRNAs was 174 (AgR015: 277523-408464), while 135 (AgB16X_g049_t01:1133282-1216863) interactions was observed to be the highest for intronic circRNAs. On average, exonic circRNAs bound to 32 miRNAs, while intronic circRNAs averaged 6 *A. suum* miRNA interactions. For human miRNA interactions with *A. suum* circRNAs, exonic circRNAs had a maximum interaction number of 2,308 (AgR015: 277523-408464), while intronic circRNAs had 1,769 (AgB16X_g049_t01:1133282-1216863). On average, exonic circRNAs had approximately 281 interactions with human miRNAs and intronic circRNAs had an average of approximately 83 interactions (Table 1). AgR015: 277523-408464 (exonic) and AgB16X_g049_t01:1133282-1216863 (intronic) were predicted to have the highest number of interactions for both worm and human miRNA interactions, suggesting that these circRNAs have a high number of binding sites located on their surface. Although there is no supporting data in the circRNA literature to explain why exonic vs. intronic circRNA binding is favorable, this could be due to the fact that exonic circRNAs contain protein coding regions, but further research into this hypothesis is needed.

**Table 1.** Summary of predicted circRNA-miRNA binding by circRNA type. Exonic circRNAs were observed to have the highest number of predicted interactions for both human and *Ascaris* miRNAs. Human miRNAs also were observed to have a significantly higher number of interactions for each of the two types of circRNAs as compared to worm miRNAs. The significant differences in exonic and intronic circRNA expression has not yet been fully established, but could be due to the ability of exonic circRNAs being translated.

| circRNA type | Highest Number of Interactions | Average number of Interactions |
| --- | --- | --- |
| Exonic circRNA |  |  |
| *A. suum* miRNAs | 174 | 32 |
| *H. sapiens* miRNAs | 2,308 | 281 |
| Intronic circRNA |  |  |
| *A. suum* miRNAs | 135 | 6 |
| *H. sapiens* miRNAs | 1,769 | 16 |

While the *A. suum* genome is not as well annotated as the human genome, it would be logical to assume that if *A. suum* circRNAs are acting as a miRNA sponge, they would have numerous miRNA interactions with *A. suum* miRNAs. Because *A. suum* circRNAs are secreted into EVs, there is also the possibility that secreted circRNAs could be acting as a miRNA sponge on human miRNAs. Comparing human and worm miRNA interactions, *A. suum* miRNA have almost 10 times less interactions than human miRNAs (Figure 7). Approximately, 9 circRNA-miRNA interactions were predicted to occur between exonic and intronic *A. suum* circRNAs and *A. suum* miRNA interactions, while *A. suum* circRNAs were predicted to interact with 115 human miRNAs, on average. Total, there were 577 circRNA – *A. suum* miRNA interactions and 645 circRNA – human miRNA interactions. 79% of circRNA – *A. suum* miRNA interactions had under 10 interactions per circRNA, while only 41% of circRNA – human miRNA interactions were under 10 interactions per circRNA. This suggests that while there is not a large difference in the number of circRNA-miRNA interactions when comparing human and worm miRNAs, the number of miRNAs per circRNA is quite different between the two species and *A. suum* circRNAs tend to have more binding sites for human miRNAs as compared to *A. suum* miRNAs.

To determine if the number of miRNA interactions was simply a reflection of circRNA length, we normalized the miRNA interaction number to length of each circRNA (Figure 8). There was not a strong linear correlation between length of *A. suum* circRNA length and *A. suum* miRNA interaction number (R^2^ = .613), suggesting that the number of *A. suum* miRNAs that an *A. suum* circRNA interacts with is not dictated by the length of circRNAs and by extension, that some *A. suum* circRNAs are explicitly enriched in miRNA interaction sites. For example, AgR030:203455-226217 (14KB) and AgR030_g080_t11: 1244134-1246599 (2KB) are both enriched by a high number of miRNA interactions (Figure 8A). Figure 8B describes a similar analysis correlating the number of human miRNA interactions with *A. suum* circRNA length. In contrast to endogenous circRNA-miRNA interactions, there does seem to be a stronger linear correlation in this relationship (R^2^: .932). This suggests that there is no specific enrichment of certain *A. suum* circRNAs for human miRNA binding.

**Figure 8.**
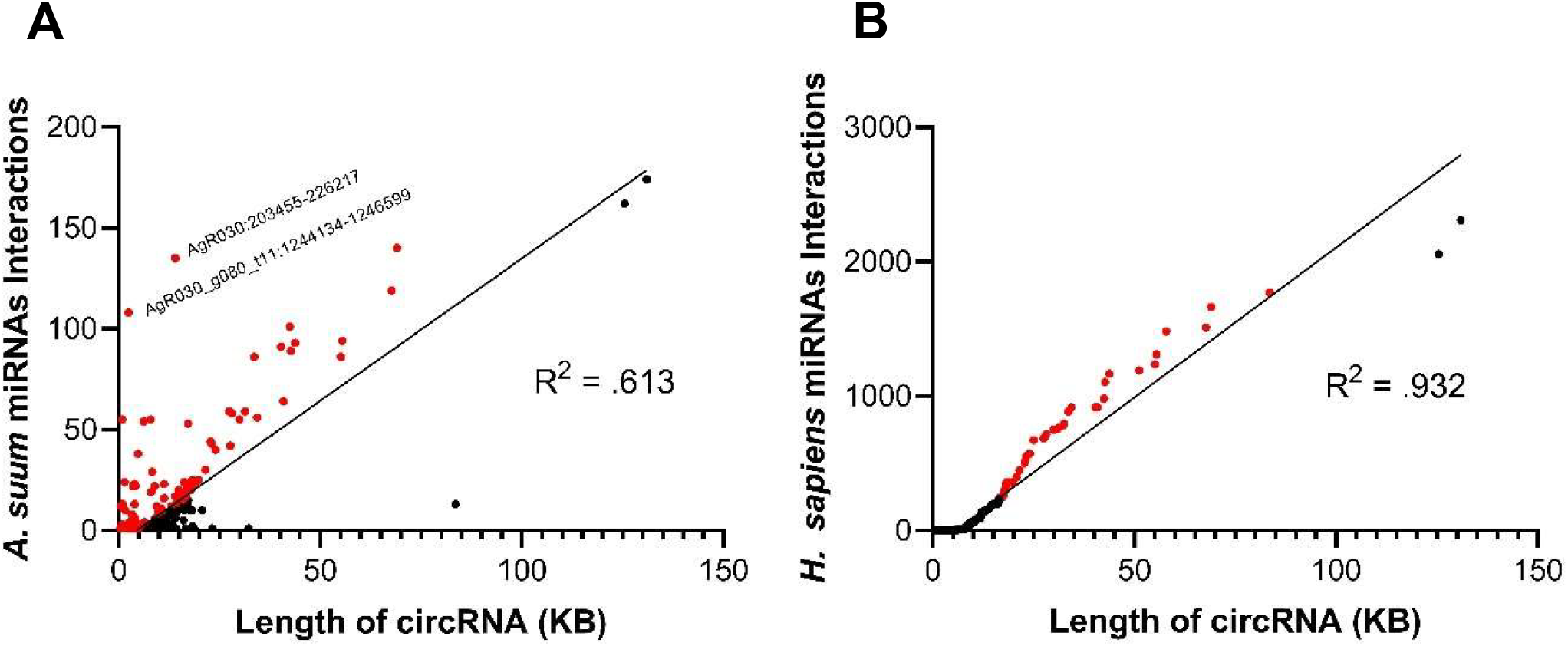
circRNA size does not influence *A. suum* miRNA binding, but host miRNA binding is related to circRNA length. To determine if the size of circRNA could be related to the amount of miRNA interactions per circRNA. Red points are located over the best line of fit and could have enriched amount of binding sites for miRNAs. **(A)** Length of circRNA per number of A. suum miRNA interactions. For A. suum miRNA binding, length does not seem to influence the number of miRNA interactions. With an R^2^ = 0.613 and a large amount of data points away from the best line of fit, this suggests *that A. suum* circRNAs could be enriched for *A. suum* miRNA binding. AgR030:203455-226217 and AgR030_g080:1244134-1246599 had a large amount of predicted *A. suum* miRNA binding compared to their relative size, suggesting they might be enriched for miRNA binding. Conversely, human miRNA binding is dependent on circRNA length **(B)**. There is a positive linear relationship observed between A. suum circRNAs and human miRNAs (R^2^= 0.932).

In addition to examining the number of miRNA interactions for each *A. suum* circRNA, we also wanted to probe the interaction from the opposite direction and determine if any *A. suum* or host miRNAs were predicated to bind to *A. suum* circRNAs. Parasite and host miRNAs with predicted circRNA interactions are summarized in Figure 9A and B, respectively.

**Figure 9.**
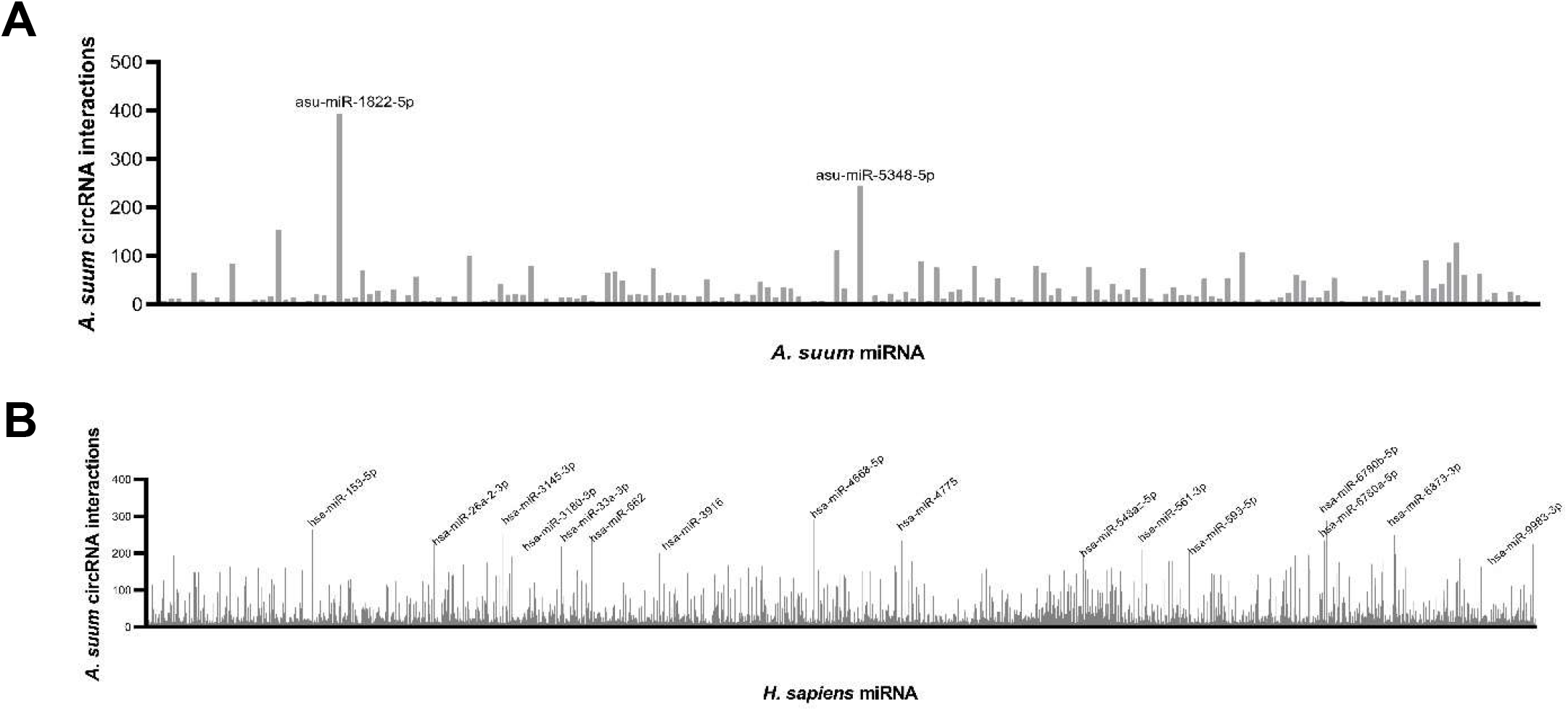
Host and *A. suum* miRNAs have multiple predicted interactions with *A. suum* circRNAs. *A. suum* **(A)** and host **(B)** miRNAs with numerous predicted interactions with circRNAs. miRNAs with less than 10 interactions were excluded from this graph. Highlighted miRNAs had over 200 interactions with circRNAs. A larger number of host miRNAs were predicted to interact with circRNAs as compared to worm miRNAs.

Again, we observed a larger number of circRNA interactions occurring with human miRNAs as compared to *A. suum* miRNAs. Total, we observed that 180 *A. suum* distinct miRNAs were predicted to interact with *A. suum* circRNAs. Two worm miRNAs had over 200 circRNA interactions, asu-miR-1822-5p and asu-miR-5348-5p (Figure 9A), but there are not any known functions or phenotypes observed with these two miRNAs. asu-miR-1822-5p did have the highest amount of circRNA interactions out of all of miRNAs at 394 total predicted interactions. Human miRNAs had a total of 2,414 predicted circRNA interactions with 16 miRNAs having over 200 circRNA interactions (Figure 9B). The majority of functions of these miRNAs are associated with cancers. The highest number of circRNA interactions for a single human miRNA was hsa-miR-4668-5p (291). hsa-miR-4668-5p has been predicted to regulate TGF-beta signaling (Bhardwaj et al., 2020). TGF-beta is an extremely important cytokine involved in the proliferation, differentiation and function of lymphocytes, macrophages, and dendritic cells (Kubiczkova et al., 2012) and dysregulation of TGF-beta, through circRNA sponging, could potentially help parasites establish infections in hosts. hsa-miR-6780b-5p had the second highest amount of predicted circRNA interactions at 289. This miRNA has been linked to insulin resistance in hepatocellular carcinoma cells (HepG2). Compared to control cells, hsa-miR-6780b-5p was upregulated in insulin resistant HepG2 cells (Li L. et al., 2020).

While this miRNA is not directly related to the immune system, there could be other processes that could be altered as a result of its sequestering by circRNA sponging that could be advantageous to parasite infection. These results suggests that secreted circRNAs could affect host gene expression through binding to miRNAs and having an impact on host immunomodulation.

To determine if circRNA could form a stable bond to worm and human miRNAs, we looked at the miRanda free energy score. A lower miRanda free energy score is related to having a more thermodynamically stable bond, and therefore, a stronger bond between circRNA and miRNA. We found that circRNAs form a more stable bond with human miRNAs as compared to *Ascaris* miRNAs (Figure 10). The free energy scores associated with human miRNAs are almost 2 times lower than *Ascaris* miRNA free energy scores. With this result, secreted circRNAs could be forming tight bonds to host miRNAs and this could lead to changes in host gene expression through secretion of *A. suum* circRNAs in EVs.

**Figure 10.**
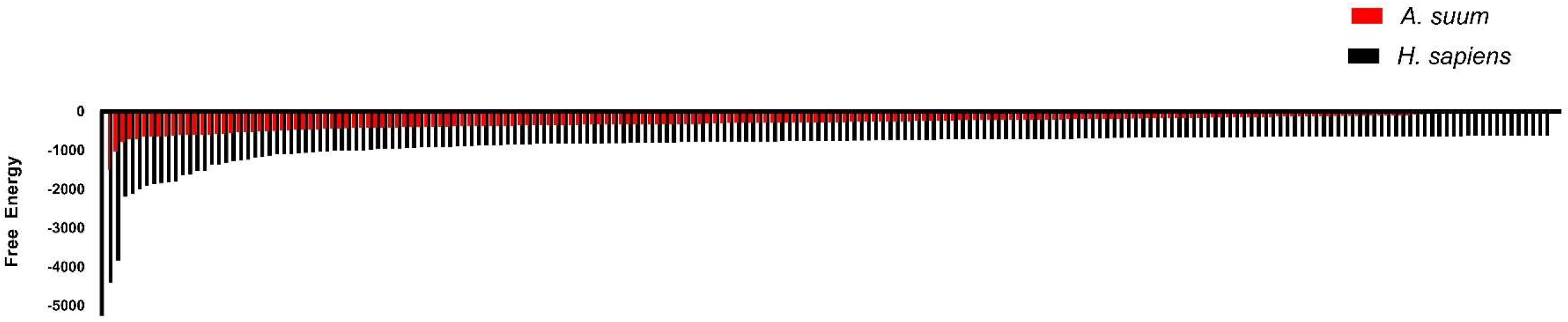
Lower miRanda free energy scores are associated with host-parasite miRNA-circRNA interactions. Lower miRanda free energy scores are associated with having a more thermodynamically stable bond occurring between circRNA and miRNA. *Ascaris* miRanda scores are seen in red, while human free energy scores are represented in black. We see that there is a more favorable bond associated between circRNAs and host miRNAs. The top 200 free energy scores from both host and worm miRNAs were graphed above.

Because miRNAs hold a vast number of functions, we wanted to ascertain if any of the host miRNAs that were predicted to bind to circRNAs had immunoregulatory functions. The percentage of immunoregulatory miRNAs was calculated and can be viewed in Table 2. Percentages were calculated for each individual miRNA out of the total predicted miRNA population. As a whole, there was not a high percentage of miRNAs that have immune regulatory functions observed, but these miRNAs could be playing important roles in host and worm biology that could be having indirect effects on the host-parasite immune interface. hsa-let-7 had the highest percentage (0.5461) and is known to influence T-cell activation and mediates cytokine expression (Gilles and Slack, 2018).

**Table 2.** Parasite circRNAs are predicted to interact with host miRNAs that have immunoregulatory functions. Frequency of each miRNA was calculated by counting the number of individual miRNAs in the sample. There was a subset of host miRNAs with predicted interactions to A. suum circRNAs that are associated with immunomodulatory functions. While known immunomodulatory miRNAs did not take up a large sum of the host miRNA demographic, miRNA with functions not directly associated with the immune system could still be affecting the host-parasite immune interface and carry beneficial functions to parasite infection.

| miRNA | Function | Frequency of miRNA in data |
| --- | --- | --- |
| hsa-let-7a-2-3p, hsa-let-7a-3p,<br>hsa-let-7a-5p, hsa-let-7b-3p, hsa-let-7b-5p,<br>hsa-let-7c-3p, hsa-let-7d-3p, hsa-let-7d-5p,<br>hsa-let-7e-3p, hsa-let-7f-1-3p, hsa-let-7f-2-3p,<br>hsa-let-7f-5p, hsa-let-7g-3p, hsa-let-7i-3p,<br>hsa-let-7i-5p | T-cell activation and mediate cytokine expression<br>(Gilles and Slack, 2018) | 0.546172705 |
| hsa-miR-10a-3p, hsa-miR-10a-5p, hsa-miR-10b-3p,<br>hsa-miR-10b-5p | T-reg cell homeostasis and inhibit Th17<br>cell functions (Chandan et al, 2020) | 0.078997548 |
| hsa-miR-124-3p, hsa-miR-124-5p | DC subset development, induction and<br>maintenance of M2 phenotype, differentiation<br>of CD4+ T-cells into Th1 cells, reduce IL-6 and<br>TNF-alpha release (Qin et. al., 2016) | 0.032688641 |
| hsa-miR-145-3p, hsa-miR-145-5p | Inhibits release of IL-6 and CXCL8<br>(Tahamtan et. al., 2018) | 0.110324162 |
| hsa-miR-146a-3p, hsa-miR-146a-5p,<br>hsa-miR-146b-3p, hsa-miR-146b-5p | Inhibition of TH1 inflammatory<br>response (Hirscheberger et al, 2018) | 0.265595206 |
| hsa-miR-150-3p, hsa-miR-150-5p | Regulates genes whose downstream products<br>encourage differentiating stem cells towards<br>becoming megakaryocytes and involved in<br>controlling B and T cell differentiation (Chandan et al, 2020) | 0.01225824 |
| hsa-miR-155-3p, hsa-miR-155-5p | Increases Th17 cell function and T effector cell<br>accumulation and action (Hirscheberger et al, 2018) | 0.044946881 |
| hsa-miR-181a-2-3p, hsa-miR-181a-3p,<br>hsa-miR-181a-5p, hsa-miR-181b-2-3p,<br>hsa-miR-181b-3p, hsa-miR-181b-5p,<br>hsa-miR-181c-5p, hsa-miR-181d-3p,<br>hsa-miR-181d-5p | Enhancement of TCR signaling and<br>phosphorylation of immunoreceptor,<br>increased M2 polarization (Hirscheberger et al, 2018) | 0.603377826 |
| hsa-miR-187-3p, hsa-miR-187-5p | Decreases IL-6 and IL-12 expression<br>(Tahamtan et. al., 2018) | 0.017706347 |
| hsa-miR-221-3p, hsa-miR-221-5p | Modulate intestinal Th17 cells (Mikami et. al., 2021) | 0.0204304 |
| hsa-miR-222-3p, hsa-miR-222-5p | Modulate intestinal Th17 cells (Mikami et. al., 2021) | 0.019068374 |
| hsa-miR-223-3p, hsa-miR-223-5p | Suppresses TLR4 signaling in macrophages and regulates<br>intestinal inflammation (Tahamtan et. al., 2018) | 0.102152002 |
| hsa-miR-24-1-5p, hsa-miR-24-2-5p, hsa-miR-24-3p | Decreases the expression of M1 markers and increases the<br>production of M2 markers (Tahamtan et. al., 2018) | 0.040860801 |
| hsa-miR-29a-3p, hsa-miR-29c-5p, hsa-miR-29c-3p,<br>hsa-miR-29b-3p, hsa-miR-29b-2-5p, hsa-miR-29b-1-5p,<br>hsa-miR-29a-5p | Helper T-cell differentiation (Liston et al 2012) | 0.476709344 |

## Discussion

While circRNAs have been described in the free-living nematode *Caenorhabditis elegans* (Memczak et al., 2013; Ivanov et al., 2015) and now a more thorough description from *Haemonchus contortus* (Zhou et al., 2021), a parasitic nematode of small ruminants, our understanding of circRNA expression and function in nematodes is lacking. A summary of circRNA biogenesis is presented in Figure 1 and the identification of exonic, intergenic and intronic circRNAs in in this study and others supports this model in nematodes.

A comparison of circRNAs in *C. elegans* with that of the two parasitic nematodes studied to date points to an overall conservation of circRNA profile. circRNA descriptions in *C. elegans* have focused on exonic circRNAs rather than intergenic or intronic circRNAs, perhaps due to the possibility of protein translation from these exonic circRNAs. A total of 1,686 exonic circRNAs have been identified in various *C. elegans* life stages (Ivanov et al., 2015; Memczack et al., 2013; Cortés-López et al., 2018). In comparison, we found a similar number of exonic circRNAs (1,178) in adult female *Ascaris suum* whilst 14,251 exonic circRNA were discovered in *H. contortus* (Zhou et al., 2021) across three life stages (the infective third stage larvae, adult male and adult female worms). The number of exonic circRNAs is comparable between *Ascaris* and *C. elegans* but significantly higher in *Haemonchus*, with the greatest number being expressed in the third stage larvae of that species. It is possible that defining the larval circRNA complement in *Ascaris* will reveal a similar level of exonic circRNA complexity. Cortés-López et al. (2018) reported that 98.2% of *C. elegans* circRNAs contained a coding sequence while 1.8% were labeled as “other”. While not explicitly stated in that manuscript, these circRNAs might be considered intronic. Compared to *A. suum* (2%) and *H. contortus* (6%), the percentage of *C. elegans* intronic circRNAs is very similar. GO terms of circRNA parental genes were only assigned for adult *C. elegans* worms with enriched pathways including organism development, determination of adult lifespan, enzyme binding and intracellular components. These GO terms are different from both *H. contortus* and *A. suum* with assigned GO terms here more focused on signaling and transcriptional pathways. This suggests that whilst overall circRNA profiles may be conserved between *C. elegans* and parasitic species, that is to say, abundance of exonic RNAs relative to intronic, the functionality of those circRNAs may be very different.

Comparing the two parasitic nematode species studied to date reveals similarities in circRNA complement. 71% of *H. contortus* circRNAs are exonic, 22% are intergenic, and 6% are intronic (Zhou et al., 2021). We identified a similar pattern in *A. suum* circRNAs, with the majority coming from exonic regions (59%), followed by intergenic (39%) and the least amount of circRNAs originating from intronic regions (2%). The majority of circRNAs in these two species are originating from protein coding regions, and may hint that an important circRNA function could be protein translation. *H. contortus* circRNA parental genes were assigned GO terms such as signaling, signal transduction, protein binding, and receptor activity. Significant GO terms assigned to *A. suum* parental genes included transcription and nucleotide binding, and whilst distinct from the *Haemonchus* assignments, still suggest that circRNAs from both worms could be functioning in important regulatory processes. *H. contortus* KEGG pathways included MAPK signaling among others. Similarly, in *Ascaris* KEGG pathways, MAPK signaling was also enriched. This suggests that pathways that circRNAs originate from in these two worms, could be performing similar functions and circRNAs could also be carrying out similar functions if they are translated. Further research into circRNAs in different tissues and *Ascaris* life stages would increase the number of circRNAs and lend to better understanding of the function of circRNAs in *A. suum*.

The majority of nematode circRNAs are derived from exonic linear RNA regions, with the possibility that these circRNAs could be translated into functional proteins. In *Drosophila*, circRNAs are known to have specific association with translating ribosomes and proteins are generated from circRNA minigenes. circRNAs also contain specific stop codons, supporting endogenous circRNA translation in *Drosophila* fly heads (Pamudurti et al., 2017). In mice, the exonic circRNA circ-ZNF609 contains a reading frame with both a start and stop codon and is associated with polysomes (Legnini et al., 2017). This circRNA is translated into a protein in a cap-independent manner, since circRNAs do not contain a 5’ cap. circ-ZNF609 was transfected into HeLa and N2A cells with two different protein isoforms produced and detected through western blot. A linearized circRNA control was also used that produced the same protein products, but was skewed towards one isoform, and was translated at two times higher magnitude than the circRNA suggesting protein translation from circRNAs may differ from linear templates. circ-ZNF609 is associated with muscular dystrophy in mice and humans and has been shown to regulate myoblast proliferation (Legnini et al., 2017). Clearly there is a precedent for exonic circRNAs to serve as substrates for protein translation and this may be an important function in parasitic nematodes.

One of the hallmark functions of circRNAs is that of a miRNA sponge. circRNAs have the ability to alter gene expression by binding miRNAs, reducing their bioavailability and leading to their loss of function. This process has been established in many organisms including humans (Panda et al., 2018), mice (Hansen et al., 2013), and *Drosophila* (Westholm et al., 2014). In humans, circRNA-miRNA sponging has been extensively studied within the context of human disease. CDR1as, the first circRNA-miRNA sponge, was discovered in mice (Hansen T. B. et al., 2013) but has also been identified in other animals, including humans where it binds miR-7 and contains over 60 binding sites for this miRNA. CDR1as has been linked to several diseases such as Alzheimer’s disease (Lukiw et al., 2013) and hepatocellular carcinoma (Yu et al., 2016) due to this sponging of miR-7. It is possible that parasitic nematode circRNAs could also function as miRNA sponges to regulate key processes in these organisms. 205 of the 1,997 circRNAs identified in adult female *A. suum* were predicted to bind to *A. suum* miRNAs. After normalizing length of circRNA to number of miRNA bindings sites, we found that *A. suum* circRNAs appeared enriched for *Ascaris* miRNA binding sites relative to their length, which may be expected if an endogenous function in this parasite is regulation of gene expression. Using the miRanda algorithm, fewer miRNAs were predicted to interact with circRNAs (194) across all three life *H. contortus* stages examined. In the sea cucumber, *A. japonicus*, the opposite was observed with 3,679 out of 3,952 circRNAs identified predicted to interact with miRNAs, while also using the miRanda algorithm (Zhao et al., 2019). Whilst these variations in predicted circRNA-miRNA interaction could be founded in differences in approach used, life stages or tissues types examined, it may also reflect differences in the functional roles for circRNAs across diverse invertebrate species.

Our laboratory and others have shown that miRNAs and other small RNA species are secreted by parasitic nematodes into the host environment via extracellular vesicles or EVs (Zamanian et al., 2015; Buck et al., 2014; Hansen E. et al., 2019; Gu et al., 2017). Delivery of those EVs to host cells elicits transcriptional changes that benefit the parasite, establishing a mechanism by which parasites can modulate host responses at the genetic level. *Heligmosomoides polygyrus* EVs inhibit genes involved in toll-like receptor signaling and IL-33 signaling (Buck et al., 2014) and also suppress macrophage activation through the IL-33 pathway, as well as driving other functionally important response pathways in those cells (Coakley et al., 2017). Filarial nematode parasites also secrete EVs that modulate macrophage phenotypes (Zamanian et al., 2015) and contain a complex miRNA cargo with explicit sequence homology to host miRNAs, suggesting parasite miRNAs could act as host miRNA mimics to affect gene expression. Relevant to this study, miRNAs found encapsulated in *A. suum* EVs are predicted to target important immune response cytokines such as IL-13, 25, and 33 (Hansen E. et al., 2019). Here, we identified *Ascaris* circRNAs in EV-enriched fractions of spent culture media, suggesting circRNAs are part of the diverse EV cargo. Further, *Ascaris* circRNAs are predicted to strongly interact with human miRNAs. We posit that parasite circRNAs could be delivered to host cells via the EVs and contribute to the transcriptional changes observed at the host-parasite interface. Supporting this hypothesis, secreted circRNAs have been shown to be functionally relevant in a wide variety of pathological settings. In colorectal cancer, circRNAs secreted via EVs have been shown to lead to drug resistance. EVs containing ciRS-122 from oxaliplatin resistant colorectal cancer cells, was delivered to drug sensitive cancer cells which then lead to resistance to oxaliplatin through sponging of miRNA-122 (Wang X. et al., 2020). In mice, circRNA circSCMH1 presence in EVs has been shown to be a biomarker for ischemic stroke (Yang et al., 2020). Lower levels of circSCMH1 in plasma correlated with a higher chance of stroke in those animals. Further, treatment with circSCMH1 improved recovery after stroke.

Defining the role circRNAs play in parasite gene regulation or manipulation of the host immune response is an important next step but will be challenging to accomplish at a technical level. Even in highly tractable model systems, circRNA functions remain elusive for this reason. In situ hybridization techniques may allow spatial localization of circRNA and miRNAs of interest in parasite tissues but whilst that might support interactions predicted *in silico*, it may fall short of providing strong functional insight. Several strategies have been used to knockdown expression of circRNAs and provide functional data. Gapmer antisense oligonucleotides can be transfected into cells or tissues of interest to drive RNaseH-mediated cleavage of circRNAs in a sequence-specific manner (Marrosu et al., Ottesen et al., 2019). Small interfering RNAs (siRNAs) have been used to good effect for downregulating circRNAs in cultured cells (Legnini et al., 2017) and may have some potential for translation to parasitic nematodes as some of these worms are susceptible to RNAi (Song et al., 2010; Verma et al., 2017). Lastly, by targeting back-splice junction sites, a CRISPR/Cas13 approach has been used to successfully screen for circRNA function (Li S. et al., 2021). This strategy may be possible if DNA transformation of parasitic nematodes becomes more feasible.

Collectively, our data shows that circRNAs are expressed in the parasitic nematode *A. suum* and are also secreted into their extracellular vesicles. These findings are in support of the recent description of *H. contortus* circRNAs by Zhou et al (2021) and better our understanding of how prarsitic nematodes may regulate gene expression. Importantly, our recognition that parasitic nematodes secrete circRNAs into the host environment is significant and adds another modality for modulation of host biology to the parasite toolkit. Clarifying the function of these circRNAs will be critical, be they as templates for protein translation or as miRNA sponges. This functional data will provide needed insight into the circRNA-miRNA-mRNA interactome, furthering our understanding of basic parasite biology but may be important for controlling these insidious pathogens. Disrupting circRNA function in the parasite or at the host-parasite interface may help prevent transmission and the establishment of infection.

## Acknowledgements

The authors would like to thank Dr. Paul Williams from the Martin Laboratory at Iowa State University for his assistance with collecting adult female *A. suum* from a local abattoir. The authors would also like to thank Drs. Ravindra Singh and Eric Ottesen at Iowa State University for providing the RNase R used in this study and for their guidance in divergent primer design.

## Disclosure of Interest

The authors report no conflict of interest.

## Supplementary Data

**Supplementary Figure 1:**
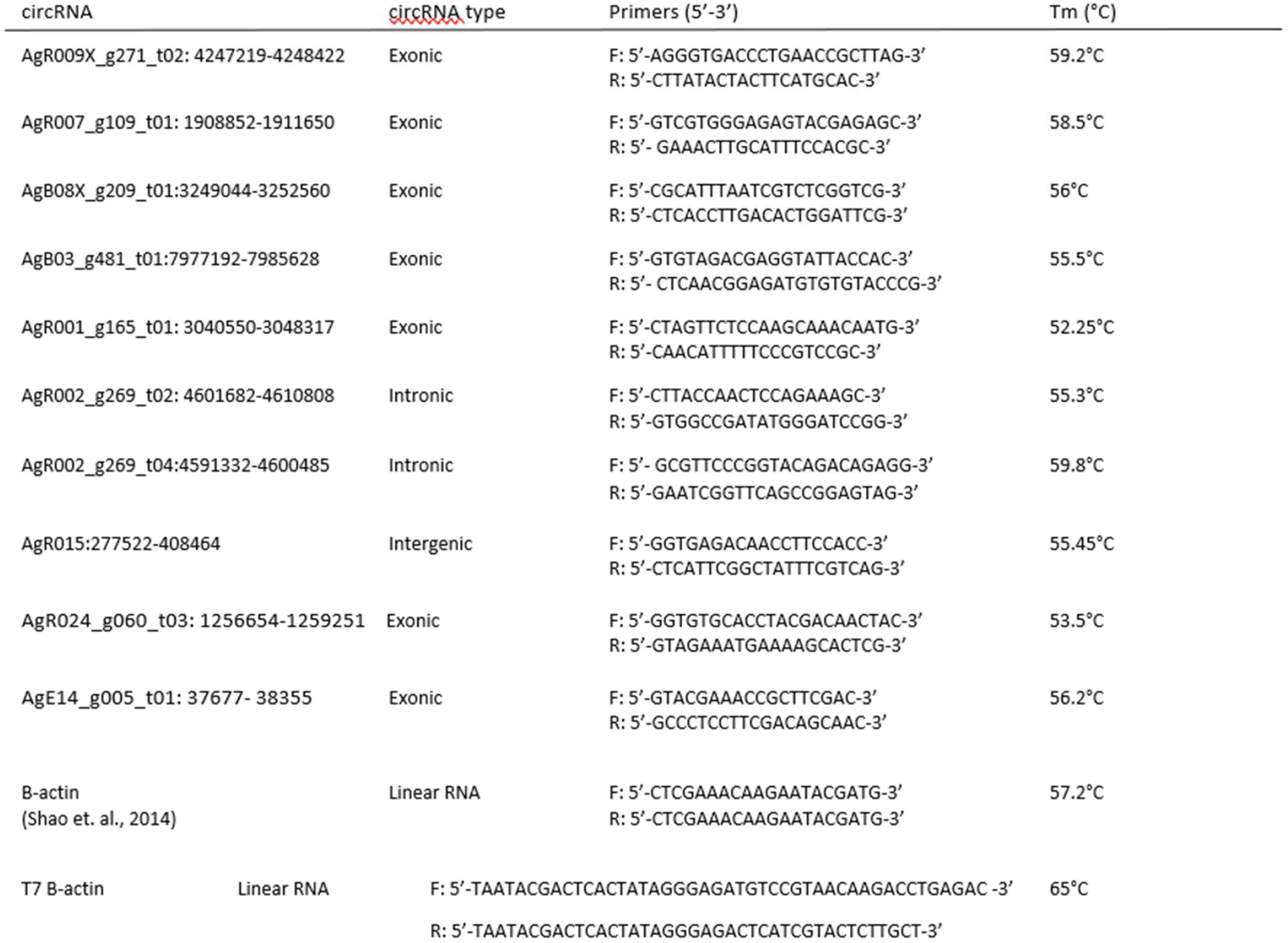
List of ovary and body wall specific circRNAs.

**Supplementary Figure 2: Primer table for primers used during qPCR validation** Primer sequence table for circRNA qPCR validation and primers for MEGAScript T7 RNA spike in.

**Supplemental Figure 3.**
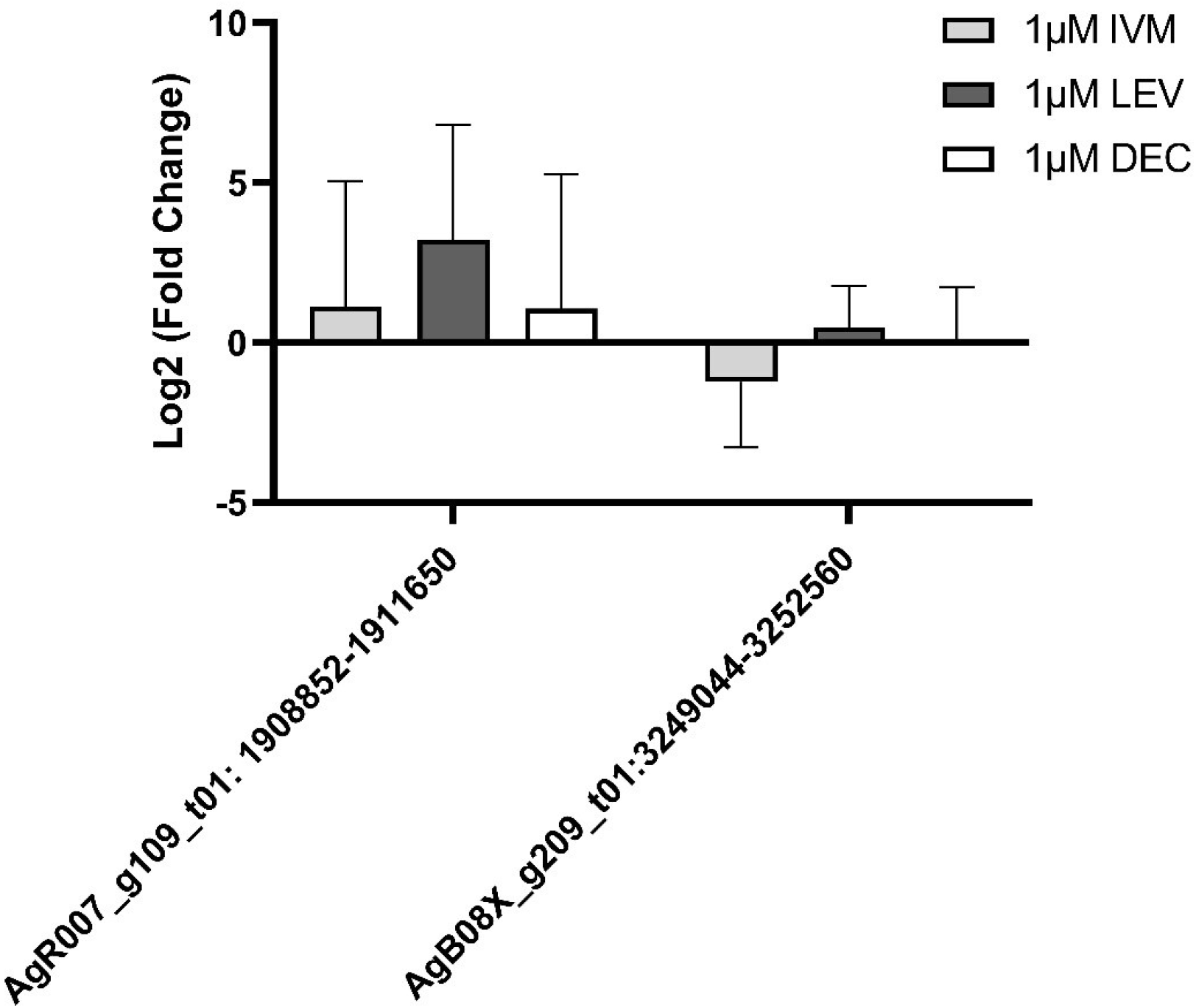
1µM IVM, DEC, and LEV do not have an effect on secreted circRNA in *A. suum* extracellular vesicles. qRT-PCR CT values were normalized to 40ng spike in RNA using 2^−ΔΔCq^ and log(2) transformation was also performed. N=3 (minimum). Mean ± SEM, p-value less than 0.05 being significant, *P<0.05, **P<0.01. Secreted circRNAs were not affected by anthelmintic treatment at 1µM drug treatment after 24 hours.

**Supplemental Figure 4.**
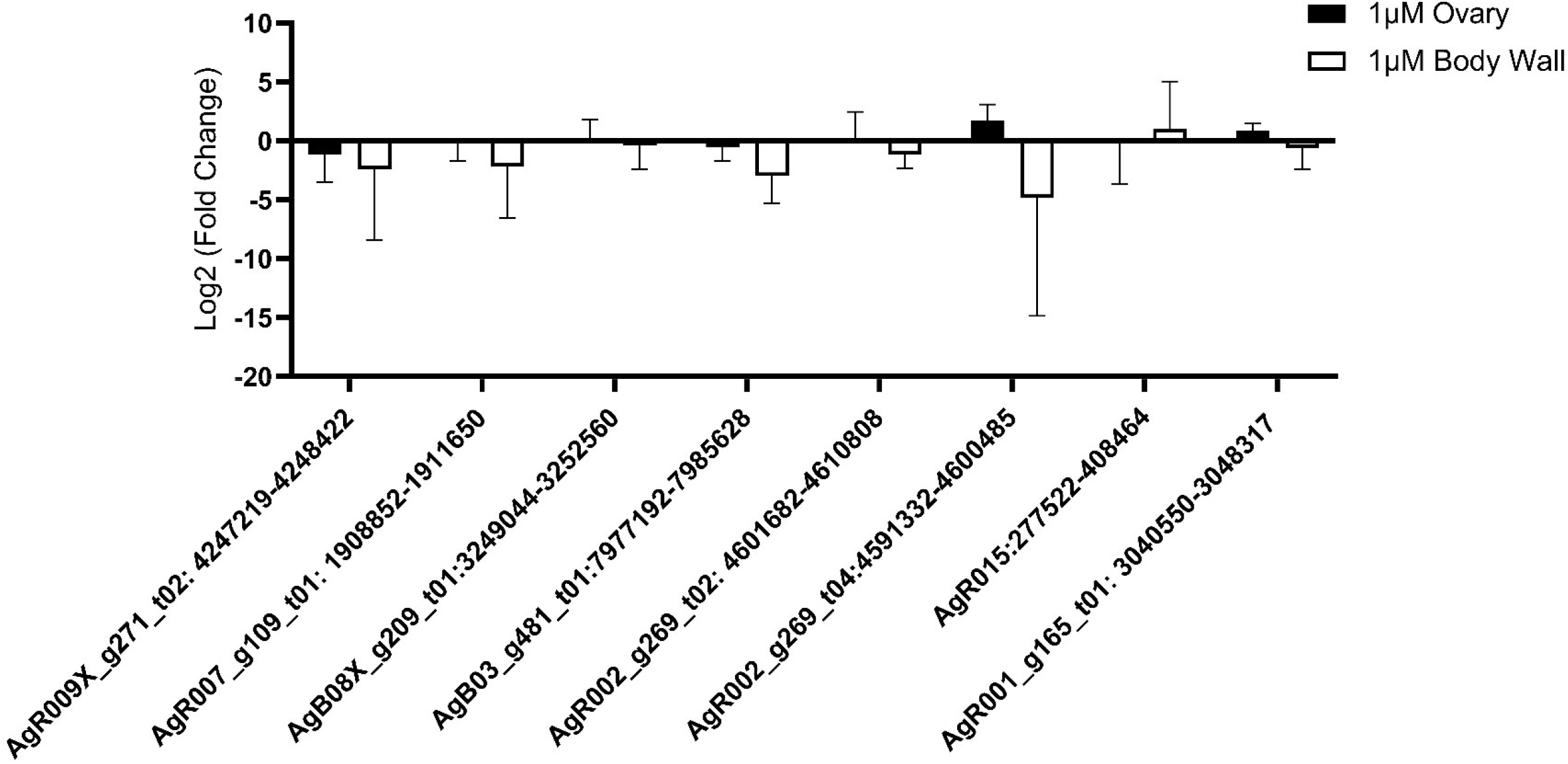
1µM IVM does not have an effect on circRNA expression in ovary and body wall tissue in *A. suum*. qRT-PCR CT values were normalized to 40ng spike in RNA using 2^−ΔΔCq^ and log(2) transformation was also performed. N=4 (minimum), Mean ± SEM, p-value less than 0.05 being significant, *P<0.05, **P<0.01. Endogenous circRNA expression was not affected by 1µM IVM treatment for either tissue type after 24-hour incubation with drug treatment.

**Supplemental Figure 5.**
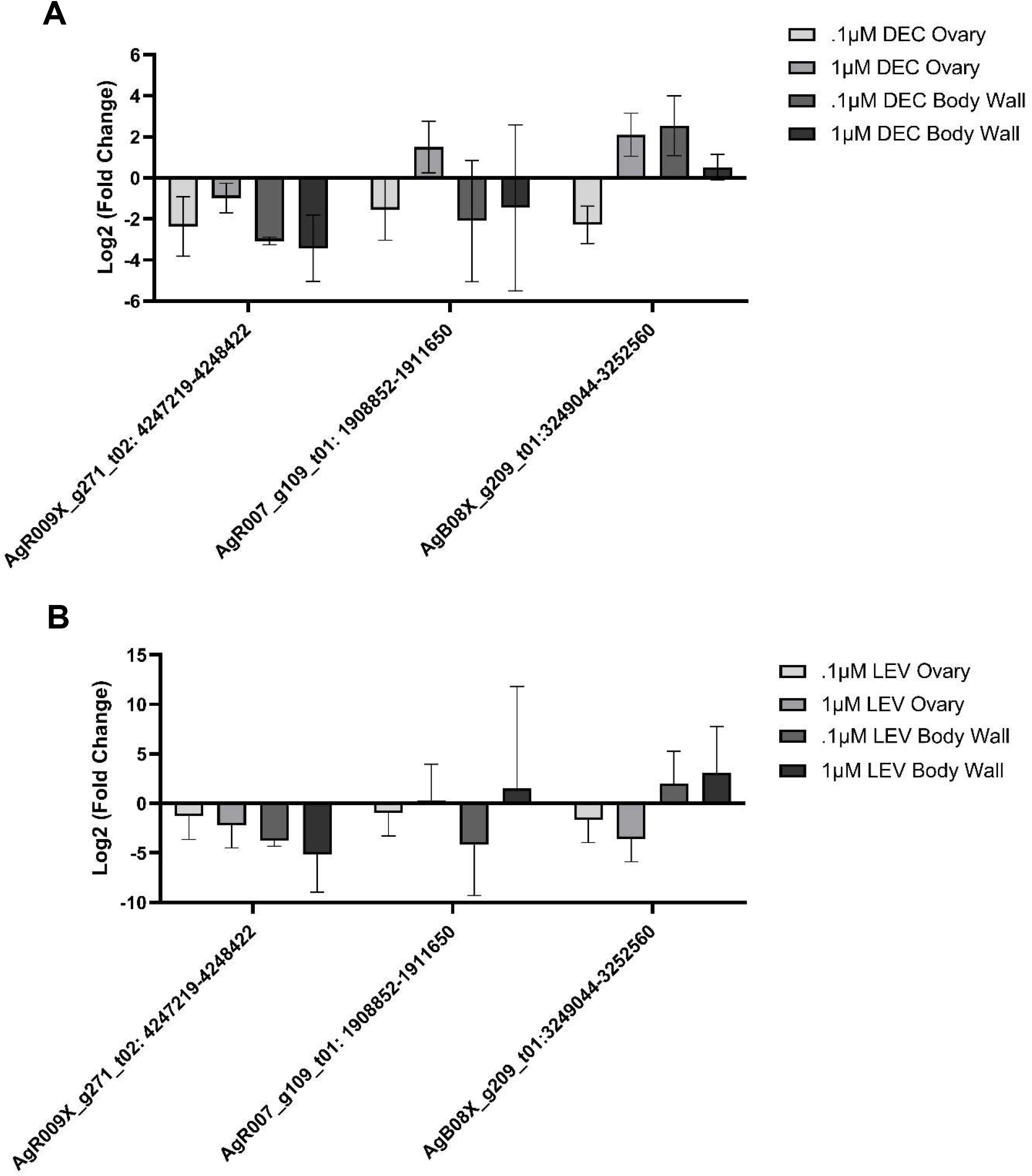
DEC and LEV do not affect circRNA expression in tissues. qRT-PCR CT values were normalized to 40ng spike in RNA using 2^−ΔΔCq^ and log(2) transformation was also performed. N=3 (minimum), Mean ± SEM, p-value less than 0.05 being significant. **(A)** circRNA expression levels DEC treated ovary and body wall at .1µM and 1µM. DEC did not have any effect on endogenous circRNA expression in ovary and body wall tissues at either concentration. Additionally, LEV treated ovary and body wall tissues did not have significant different in circRNA expression level compared to control at either concentration **(B)**.

**Supplementary Figure 6:** Notes and scripts used to produce expression analysis are available at https://github.com/ISUgenomics/Kimber_CircRNA

**Supplementary Figure 7:** Raw data from sequencing can be viewed using bio-project number <u>PRJNA750737</u> with SRA numbers <u>SRR15295818</u> - <u>SRR15295823</u>.

## Notes

https://github.com/ISUgenomics/Kimber_CircRNA

https://dataview.ncbi.nlm.nih.gov/object/PRJNA750737

https://dataview.ncbi.nlm.nih.gov/object/SRR15295818

https://dataview.ncbi.nlm.nih.gov/object/SRR15295823

### Competing Interest Statement

The authors have declared no competing interest.

